# Frizzled2 receives the WntA morphogen during butterfly wing pattern formation

**DOI:** 10.1101/2023.04.11.536469

**Authors:** Joseph J Hanly, Ling S Loh, Anyi Mazo-Vargas, Teomie S Rivera-Miranda, Luca Livraghi, Amruta Tendolkar, Christopher R Day, Neringa Liutikaite, Emily A Earls, Olaf BWH Corning, Natalie D’Souza, José J Hermina-Perez, Caroline Mehta, Julia Ainsworth, Matteo Rossi, W. Owen McMillan, Michael W Perry, Arnaud Martin

**Affiliations:** Department of Biological Sciences, The George Washington University, Washington, DC, USA; Smithsonian Tropical Research Institute, Gamboa, Panama; Department of Entomology, Purdue University, West Lafayette, IN, USA; Department of Zoology, University of Cambridge, Cambridge, UK; Epigenetics and Stem Cell Biology Laboratory, National Institute of Environmental Health Sciences, Durham, USA; Department of Cell and Developmental Biology, UC San Diego, La Jolla, CA, USA; Division of Evolutionary Biology, Ludwig Maximilian University, Munich, Germany

**Keywords:** evo-devo, pattern formation, wing epithelium, Wnt signaling, Frizzled receptors, morphogens, planar cell polarity, decoy receptor

## Abstract

Butterfly color patterns provide visible and biodiverse phenotypic readouts of the patterning processes that occur in a developing epithelium. While the secreted ligand WntA was shown to instruct the color pattern formation in butterflies, its modes of reception and signal transduction remain elusive. Butterfly genomes encode four homologues of the Frizzled-family of Wnt receptors. Here we show that CRISPR mosaic knock-outs of *frizzled2* (*fz2*) phenocopy the color pattern effects of *WntA* loss-of-function in multiple nymphalids. While *WntA* mosaic clones result in intermediate patterns of reduced size, consistently with a morphogen function, *fz2* clones are cell-autonomous. Shifts in pupal expression in *WntA* crispants show that *WntA* and *fz2* are under positive and negative feedback, respectively. Fz1 is required for Wnt-independent planar cell polarity (PCP) in the wing epithelium. Fz3 and Fz4 show phenotypes consistent with Wnt competitive-antagonist functions in vein formation (Fz3 and Fz4), wing margin specification (Fz3), and color patterning in the Discalis and Marginal Band Systems (Fz4). Overall, these data show that the WntA/Frizzled2 morphogen-receptor pair forms a signaling axis that instructs butterfly color patterning, and shed light on the functional diversity of insect Frizzled receptors.

## Introduction

The Wnt-family ligand WntA instructs color pattern formation during butterfly wing development and establishes the position and limits of color fields across Nymphalidae (Martin and Reed, 2014; Mazo-Vargas et al., 2017). Several lines of evidence pinpoint WntA signaling as a driver of color pattern evolution. First, regulatory shifts of *WntA* spatial expression have repeatedly driven adaptive variations, as evidenced by the identification to date of more than 20 *WntA* alleles associated with wing mimicry in variable species of the *Heliconius*, *Limenitis,* and *Elymnias* butterfly radiations (Gallant et al., 2014; Martin et al., 2012; Morris et al., 2020; Ruttenberg et al., 2021; Van Belleghem et al., 2017). In these systems, *WntA* alleles explain pattern differences between populations and morphs of the same species. Additionally, mapping, expression, and loss-of-function experiments show that *WntA* repeatedly shaped the spectacular ressemblance of *Heliconius* co-mimics that converged onto identical patterns, in spite of historical drift in their patterning systems (Concha et al., 2019; Van Belleghem et al., 2020). Finally, while *WntA* regulation and function appear largely conserved across butterflies that follow a bauplan-like organization of wing patterns known as the Nymphalid Ground Plan (NGP), it has also evolved atypical modes of expression associated with novel pattern elements, such as in the control of white-orange intravenous markings in the monarch butterfly (Mazo-Vargas et al., 2017; Mazo-Vargas et al., 2022). This widespread genetic parallelism (Scotland, 2011) suggests that *WntA* is a favored genomic locus for wing pattern variation (Martin and Courtier-Orgogozo, 2017; Van Belleghem et al., 2021). A high density of open-chromatin elements at this locus, each involved in the regulatory control of pattern phenotypes may underlie the apparent evolvability of *WntA* expression in nymphalids (Mazo-Vargas et al., 2022).

In addition to its insights on genetic evolution, *WntA* wing patterning also brings the opportunity to study morphogenetic signaling in a novel developmental system, where color states provide a direct phenotypic readout of positional information during development. While *WntA* evolved at the base of Metazoa (Darras et al., 2018; Kraus et al., 2016; Somorjai et al., 2018), it was lost in vertebrates as well as in Cyclorrhapha, a fly lineage that includes *Drosophila* (Hanly et al., 2021), making it a particularly understudied Wnt ligand. We previously reported that CRISPR mosaic knock-outs of *Wntless* (*Wls*) and *porcupine* (*por*) phenocopy *WntA* knock-outs, which implies that WntA is a secreted lipid-modified ligand that is processed via the classical Wnt secretory pathway (Hanly et al., 2021). In addition to its role in Wnt maturation and secretion, the fatty acid thumb added by Por-dependent acylation is essential for steric binding of Wnt ligands into the Cysteine Rich Domains (CRD) of receptors of the Frizzled family (Alvarez-Rodrigo et al., 2023; Hoppler and Moon, 2014; Nile and Hannoush, 2019; Povelones and Nusse, 2005). Studies of Wingless (Wg; ortholog of Wnt1) in *Drosophila* development have shown that Frizzled1 (Fz, here abbreviated Fz1) and Frizzled2 (Fz2) mediate short-range Wnt reception with different efficiencies, and that they are transcriptionally repressed by Wg signaling (Bhanot et al., 1999; Chen and Struhl, 1999; Moline et al., 2000). In the *Drosophila* wing disc, Fz1 is the sole Frizzled receptor necessary for the Wnt-independent Fz/Planar Cell Polarity (Fz/PCP) pathway (Ewen-Campen et al., 2020); Fz2 maintains long-range activation of the Wnt pathway in cells that are beyond the reach of extracellular Wg (Chaudhary et al., 2019); Fz3 is a decoy receptor of Wg that attenuates signaling (Sato et al., 1999; Schilling et al., 2014); and Fz4 has no known function in wings (Ewen-Campen et al., 2020). In butterflies, an RNAseq study showed that the butterfly homolog of *frizzled2* (*fz2*) is differentially expressed across color regions during early pupal wing development (Hanly et al., 2019).

To further shed light on the mechanisms of *WntA* reception, here we investigated functions of the Frizzled family seven-transmembrane domain receptors during butterfly wing development. Using CRISPR-mediated mutagenesis across several butterfly species of the Nymphalidae family, we show that *fz2* is the only Frizzled-family receptor required for the formation of WntA-dependent patterns. In contrast, *frizzled1* mosaic knock-outs indicate it mediates a conserved Fz-PCP pathway involved in scale orientation. Finally, we show that loss-of-function of *frizzled3* (*fz3*) and *frizzled4* (*fz4*) do not affect *WntA* patterns, but that these two receptors may play negative roles on WntA-independent Wnt signaling during wing development.

## Results

### Nymphalid butterfly genomes encode four Frizzled receptors

Frizzled receptors are seven-transmembrane domain proteins belonging to the G protein-coupled receptor family. Most non-vertebrate genomes encode four Frizzled orthology groups that originated before the Bilateria/Cnidaria split, including the four receptors annotated in *Drosophila* (Janssen et al., 2015; Schenkelaars et al., 2015). We recovered orthologues of these four genes in nymphalid butterfly genomes, hereafter numbered by orthology groups and abbreviated *fz1*, *fz2*, *fz3* and *fz4* (Fig. S1A, **Supplementary File 1**). Both *fz1* and *fz2* have a conserved KTxxxW motif (Fig. S1B), required for the recruitment of Dishevelled (Dsh), which relays two pathways in *Drosophila*, Fz1/2-dependent canonical Wnt signaling, and Fz1/PCP signaling (Ewen-Campen et al., 2020; Gao and Chen, 2010; Mieszczanek et al., 2022; Tauriello et al., 2012). In addition, the KTxxxW is invariable between Fz2 orthologues between *Drosophila* and Lepidoptera, showing a conserved KTLESW peptidic chain thought to contain a glutamyl-endopeptidase site necessary for the Frizzled nuclear import (FNI) pathway in *Drosophila* (Mathew et al., 2005; Restrepo et al., 2022). In contrast the KTxxxW domains are degenerated in both Fz3 and Fz4 in Diptera and Lepidoptera. Overall, amino-acid sequences from the cytoplasmic domains suggest that nymphalid Frizzled receptors may function similarly to the dipteran model *Drosophila*, with Fz1/2 signaling via Dishevelled recruitment, Fz2 potentially subject to the FNI Wnt pathway, and Fz3/4 playing alternate roles. *In situ* hybridizations of the four *fz* orthologues of *V. cardui* show these four genes are expressed in wing imaginal disks from fifth instar larvae, but, without an apparent association with color patterns, unlike *WntA* at this stage (Fig. S1C-G). These data suggest functional divergences in the use of Frizzled receptors in butterfly wing development.

### Fz2 is required for WntA patterning across nymphalid butterflies

A previous RNAseq analysis in *Heliconius* pupal wings suggested spatial complementarity of *fz2* expression levels with *WntA* (Hanly et al., 2019). We thus sought to test the function of *fz2*, and used CRISPR somatic mutagenesis to generate mosaic knockouts (mKOs) of *fz2* in a total of 6 species previously targeted for *WntA* mutagenesis (Concha et al., 2019; Mazo-Vargas et al., 2017; Mazo-Vargas et al., 2022).

In all instances, *fz2* mKOs phenocopied the effects of WntA loss-of-function, with the removal or shifting of specific patterns on all four wing surfaces. Specifically, the phenotypes observed in *Vanessa cardui* and *Junonia coenia* show (*i*) complete losses of the Central Symmetry System (CSS) – a conserved pattern element that expresses *WntA* (Martin and Reed, 2014; Mazo-Vargas et al., 2017); (*ii*) shifts in peripheral patterns; and (*iii*) specific reductions or loss of other WntA-expressing patterns such as the *Vanessa* forewing eyespots and the *Junonia* Basalis orange-black pattern (Figs. 1A,C). In *Agraulis incarnata*, *fz2* crispants reproduce all losses of silver spots known to express *WntA* in fifth instar wing disks, as well as an expansion of silver domains in the anterior hindwing, a phenomenon also observed in WntA mKOs (Martin and Reed, 2014; Mazo-Vargas et al., 2017). We knocked out *fz2* in a pair of *Heliconius* butterfly co-mimics that convergently evolved a red forewing band. The *fz2* mKOs recapitulated the species-specific effects of *WntA* removal (Concha et al., 2019; McMillan et al., 2020): a subtle proximal extension of the red band revealing a cryptic yellow dot in *Heliconius melpomene rosina*, versus an extensive proximal extension in *Heliconius erato demophoon*. Additionally, we repeated *WntA* mKOs in the monarch butterfly *Danaus plexippus* (Fig. S2), and obtained more extensive phenotypes than previously reported, reinforcing that *WntA* evolved a role in antagonizing the formation of white patterns that outline black vein markings and as discal and marginal spots (Mazo-Vargas et al., 2017). Loss of function of *fz2* reproduced these effects, with white patterns expanding in these fields.

**Figure 1.**
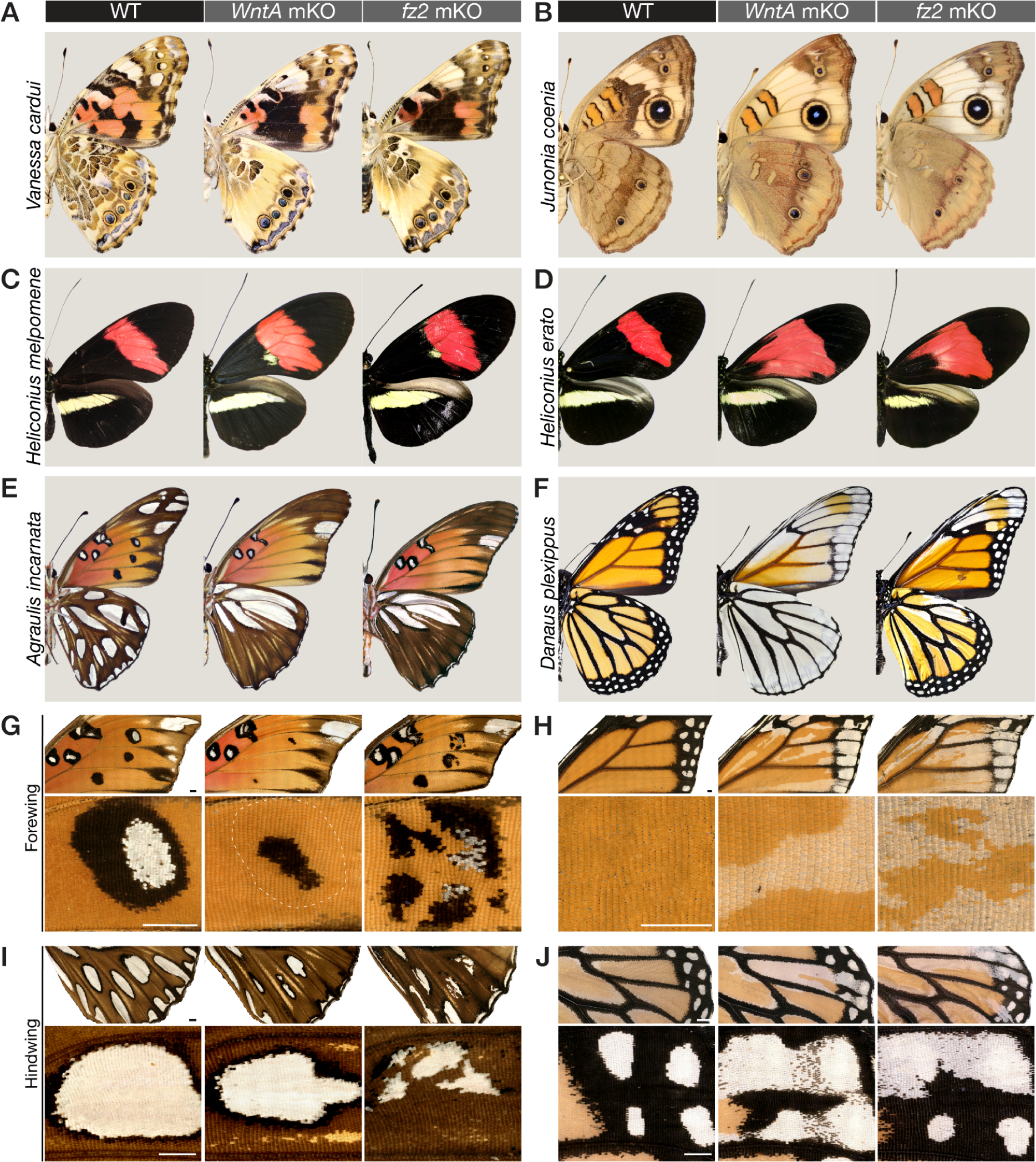
CRISPR-induced *fz2* mKOs phenocopy *WntA* mKOs in six nymphalid species, but in a cell-autonomous rather than non-autonomous fashion. **(A-F)** Ventral views comparing the effects of *WntA* and *fz2* mKOs in individuals with extensive phenotypes – *ie.* with no or little mosaicism – except in monarchs where no fully penetrant *fz2* phenotype was obtained (F, right panel). (**G-J)** Magnified views of mutant clones in individuals with high-mosaicism, contrasting cell non-autonomy of *WntA* mKOs (rounded boundaries), to cell-autonomy for *fz2* (jagged boundaries with conservation of pattern layers). (**G-H**) Ventral forewings with magnified views of the M_3_-Cu_2_ region in *A. incarnata* (G) and Cu_2_-Cu_1_ in *D. plexippus* (H). (**I-J**) Ventral hindwings with magnified views of the M_3_-Cu_2_ region in *A. incarnata* (I) and Cu_2_-Cu_1_ in *D. plexippus* (J). Scale bars = 1 mm.

As observed in *WntA* mKOs, *fz2* mKOs by injection of a Cas9/sgRNA duplex in syncytial embryos resulted in healthy adult G_0_ butterflies without detectable deleterious impacts on wing development (Figs. S3-6; Table S1). In particular, we did not observe the high rates of missing wings observed with *por* and *wls* mKOs (Hanly et al., 2021). This suggests *fz2* alone is dispensable for mediating Wnt functions that are essential for normal wing growth akin to perturbation assays in *Tribolium* embryos and *Drosophila* wings, where *fz2* loss-of-function effects are phenotypically silent due to functional redundancy with *fz1* (Beermann et al., 2011; Ewen-Campen et al., 2020). However, *fz2* crispants showed highly efficient and penetrant wing color pattern phenotypes in all tested species (Fig. 1).

A closer look at *WntA* and *fz2* crispants reveals interesting differences in the shape and aspect of mosaic clones (Figs. 1G-J). *WntA c*lones always appear rounded: this is consistent with the expected cell non-autonomous effects of a paracrine signaling ligand. In contrast, *fz2* clones show elongated, jagged shapes in the proximo-distal direction, consistent with expectations for cell autonomous effects, and reminiscent of the shape of butterfly wing clones obtained with mKOs of selector transcription factors and pigment pathway genes (Livraghi et al., 2017; Tunström et al., 2023; Westerman et al., 2018; Zhang et al., 2017a; Zhang et al., 2017b), which we would also expect to be cell autonomous. Taken together, these data show that Fz2 reception is necessary for mediating WntA patterning, and that WntA signals reach several cell diameters away from their source.

### WntA signaling represses *fz2* expression in pupae

Together, the WntA ligand and Fz2 receptor form a paracrine signaling axis required for the induction of CSS patterns, as well as for the proper positioning of pattern elements close to the wing margin, such as the distal parafocal elements (dPF), Marginal Band System (MBS), and the WntA-dependent forewing Border Ocelli (fBOc) in *Vanessa* (Figs. 2A, S7 and S8A-C). *In situ* hybridizations in a number of species had previously shown that *WntA* expression already prefigures pattern elements in late larval wings (Jiggins et al., 2017; Martin and Reed, 2014; Mazo-Vargas et al., 2017). However, current evidence suggests that extracellular WntA signaling mostly occurs after the onset of metamorphosis. Indeed, injections of heparin, an extracellular Wnt interactor that result in Wnt-dependent pattern expansions (Martin and Reed, 2014; Mazo-Vargas et al., 2017; Sourakov, 2018; Sourakov and Shirai, 2020), are effective within 15% of pupal development – typically within 24 hr after pupa formation (APF) at optimal rearing temperatures. Further, scale organ precursor cell differentiation also occurs around 13-15% (Dinwiddie et al., 2014), and the competency of tissue grafts to induce ectopic patterns (Nijhout, 1991), or developmental studies of eyespot formation (Brunetti et al., 2001; Monteiro, 2015), all imply the existence of paracrine signals active during early pupal development. We thus hypothesized that extracellular WntA signaling function actually occurs after the onset of metamorphosis.

**Figure 2.**
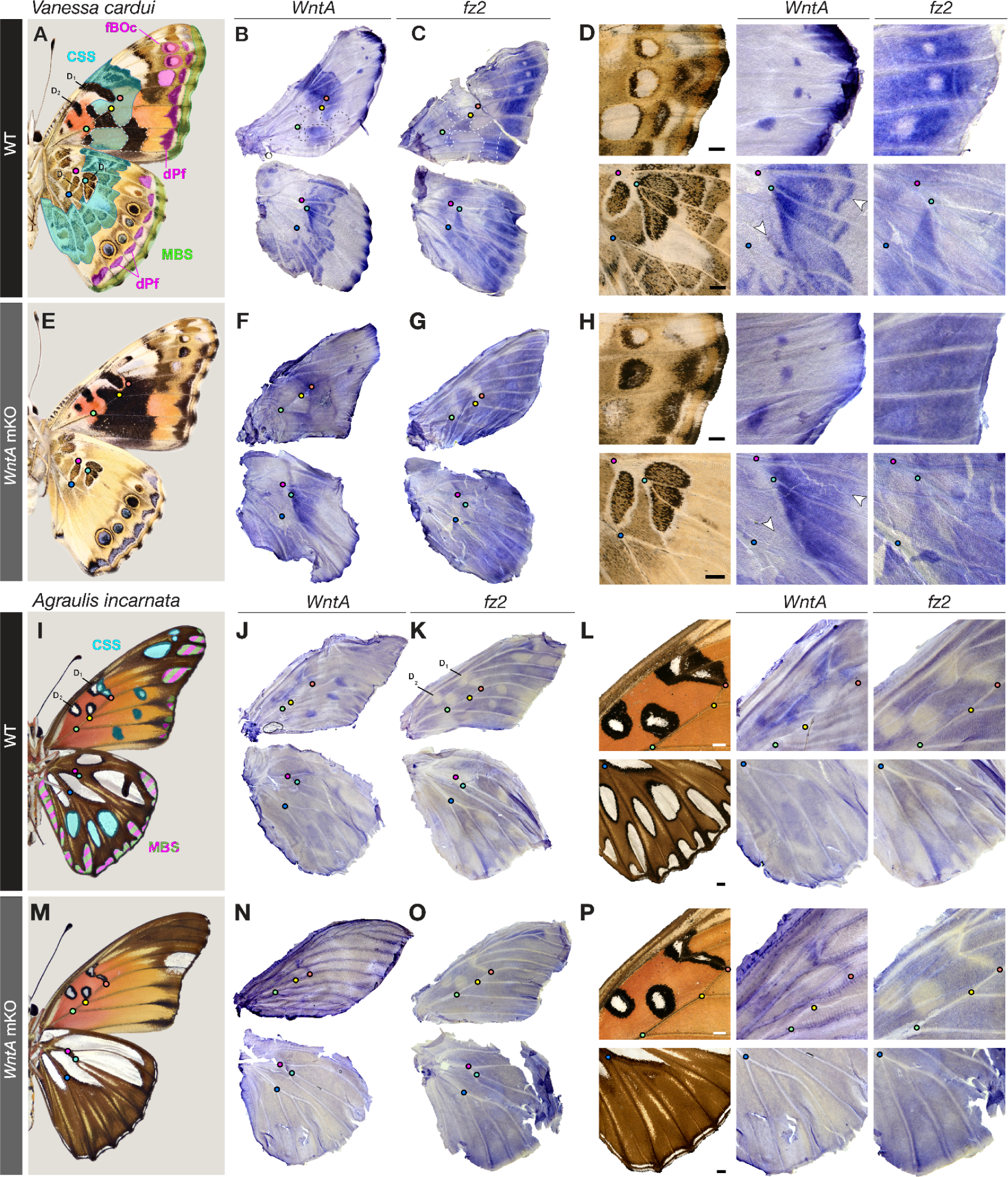
Expression of *WntA* and *fz2* in pupal wings is under the control of positive and negative feedback. **(A)** Ventral *V. cardui* annotated with its main pattern homologies from the Nymphalid groundplan (NGP): Discalis elements (D_1_ and D_2_); CSS: Central Symmetry System (cyan); BoSS: Border Ocelli Symmetry System (magenta); fBOc: forewing Border Ocelli; dPf: distal Parafocal elements; MBS: Marginal Band System (green). Color dots mark vein intersection landmarks (red: crossvein-M_3_; yellow: M_3_-Cu_1_; green: Cu_1_-Cu_2_). **(B-D)** *In situ* hybridizations (ISH) for *WntA* and *fz2* mRNA in WT *V. cardui* wings, and magnified views in the anterior tip of the forewing (top) and median hindwing (bottom). (**E-H**) Equivalent experiments to (A-D) following *WntA* knock-out. **(I)** Derivation of the NGP in ventral *A. incarnata*. The CSS is dislocated, and marginal patterns may include partial homology with dPF elements (green and magenta). **(J-L)** *In situ* hybridizations (ISH) for *WntA* and *fz2* mRNA in WT *A. incarnata* wings, and magnified views in the anterior tip of the median forewing and hindwing. (**M-P**). Expression assays (similar to J-L) following *WntA* knock-out. Comparisons of *WntA vs fz2* in *V. cardui* (B-D, F-H) are shown in contralateral tissues from the same individual. Scale bars = 1 mm.

We examined *WntA* and *fz2* mRNA expression in early pupal wings in order to gain further insights into WntA/Fz2 patterning (Figs. 2, S8). Consistent with an instructive role for pattern induction in pupal wings, *WntA* expression precisely prefigured the position and shape of WntA-dependent pattern elements, including delineations of the adult patterns and graded expression levels that prefigure their textural details in a more highly correlated way than in larvae, in all species. (*e.g.* Figs. 2D, S8F). On the other hand, *fz2* is generally expressed in an anticorrelated pattern with *WntA*. In *Agraulis*, as in *Vanessa*, we found a complementary expression of *fz2* to *WntA* in pupal stages in WT wings (Fig. 2J-L). In summary, *fz2* showed a complementary expression to *WntA,* with low staining in *WntA*-positive regions across all the WT tissues we assayed. This is indicative of WntA-dependent repression of *fz2* expression in the nymphalids, a phenomenon observed with Wg and *fz2* in larval *D. melanogaster* wing discs (Cadigan et al., 1998; Zhang and Carthew, 1998).

In order to test if WntA represses expression of *fz2*, we performed *fz2 in situ* hybridizations in *WntA* G_0_ mosaic knockouts of *V. cardui* and *A. incarnata.* In both species, patterns of *fz2* expression were markedly affected by *WntA* KO, and all WntA-expressing patterns lost the local repression of *fz2* (Figs. 2E-H, S8F’-G’). The local repression of *fz2* requires a functional WntA. Thus, while Fz2 is a positive regulator of WntA signaling, *fz2* expression is also repressed in the presence of WntA signaling, and this mechanism is likely to enforce a transcriptional negative feedback loop on this pathway.

### Fz2 mediates the patterning functions of other Wnts

The mRNA signal for *fz2* remained low in D_1_ and D_2_ Discalis patterns in the WntA-deficient wings of the two species we assayed (Fig. 2K, L, O **and** Fig. S8H’, I’), implying that a WntA-independent signal repressed *fz2* in these patterns. In parallel, a closer examination of these phenotypes reveal that *fz2* KOs consistently induced black pattern expansions in *Vanessa* forewing D_1_ and D_2_ (Fig. S9A). This effect is not detected in *WntA* KOs, implying that Fz2 receives additional patterning signals from other ligands in this part of the wing. In the *Vanessa* forewing, this may include Wg, as D_1_ and D_2_ express both *wg* and *WntA* (Fig. S9A-G). We infer that fz2 mediates the patterning functions of Wg or another Wnt, and is under additional negative transcriptional feedback from this input, similarly to WntA in other color patterns, or to the Wg/Fz2 pair in *Drosophila* (Cadigan et al., 1998; Chaudhary et al., 2019; Schilling et al., 2014). On the other hand, in the two nymphalids *Agraulis* and *Junonia* (Figs. S9H-I and S10), *wg*-positive patterns such as the D_2_ element are unaffected by the *fz2* KO, suggesting that if *fz2* is necessary for Wg color patterning, it must be redundantly with another receptor in certain contexts or species.

### WntA signaling feedback refines *WntA* expression

In addition to the transcriptional negative feedback on *fz2*, our data also suggest that *WntA* expression is under the control of positive feedback. This effect is most visible in the CSS patterns of *Vanessa*, where the spatial complexity of *WntA* pupal expression disappears when WntA signaling is removed. For instance, *WntA* expression in WT hindwings prefigures textural details of the CSS, with sharp outer boundaries, and a complex composition of undulating patterns in between (Fig. 2D’). These features of *WntA* expression are lost upon *WntA* KO (Fig. 2H’), meaning that a positive feedback of WntA signaling likely refines the sharply delineated and undulatory aspect of the ventral hindwing CSS in *Vanessa*. Similarly, in the WT posterior forewing, *WntA* is expressed in wide blocks that mark the presumptive orange patterns (Fig. S2G). In WntA-deficient pupal forewings, *ie.* in the absence of a WntA positive feedback loop, *WntA* expression is depleted from the inside of these patterns but remains high in two continuous, lateral lines (arrowheads in Fig. S2G).

Elsewhere, *WntA in situ* hybridizations show reduced staining following *WntA* mKO in *Vanessa* and *Agraulis* marginal patterns as well as in the *Agraulis* CSS (Figs. 2, S8) indicating the existence of feedback mechanisms. As an exception, *WntA* expression in the *Vanessa* forewing eyespots appears unchanged upon mKO and may be largely WntA-independent (Fig. 2H). In summary, the shifts in *WntA* expression observed in loss-of-function crispants point at a positive feedback necessary for the elaboration of CSS features in *Vanessa* wings. The importance of this feedback may vary across regions and species.

### WntA/Fz2 signaling provides proximodistal positioning to peripheral patterns

The patterns closest to the wing margin consist of stripe patterns that form the Marginal Band System (MBS) as well as the distal Parafocal element (dPf), a pattern often taking the shape of individuated arcs or chevrons between each wing vein compartment (Nijhout, 2017; Otaki, 2012; Taira et al., 2015). *WntA* is strongly expressed at the vein tips of *V. cardui* pupal hindwings (Fig. 3A). Transcripts of *fz2* are repressed across a larger domain across the periphery, suggesting local repression by negative feedback from marginal signals (Fig. 3B). *WntA* mosaic KOs leave border ocelli (eyespots) and MBS patterns unaffected in this system (Hanly et al., 2021; Mazo-Vargas et al., 2017), but result in a visible shift and reversal of the dPf blue chevron shapes (Fig. 3C-D). These phenotypes are closely reproduced in *fz2* crispant butterflies, with dPF elements occurring a distalization and thickening (Fig. 3E). Unlike WntA continuous clonal effects of dPf elements, clonal mosaics of *fz2* KO result in seemingly cell-autonomous gains of blue scales (Fig. 3D’, E’). Similar effects are visible in monarchs (Fig. 1H), where WntA/Fz2 are required for establishing the black contours of marginal white spots. Together, these expression and loss-of-function data suggest that *WntA* diffuses away from the vein tips and requires *fz2* for proper positioning and shaping of the dPF elements.

**Figure 3.**
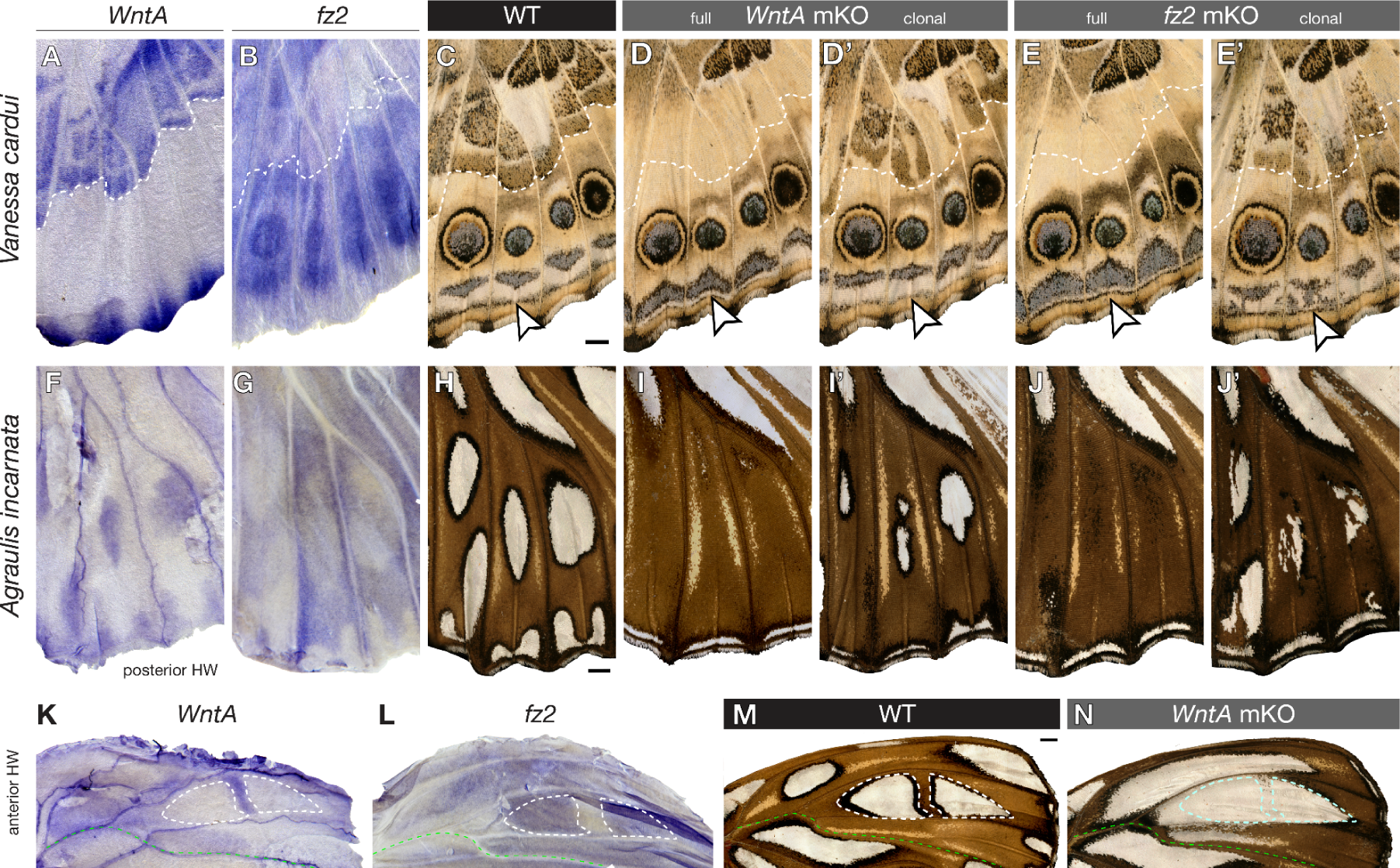
WntA/Fz2 signaling in the formation of ventral hindwing patterns. (**A-B**) In *V. cardui* pupal wing mRNA stainings, *fz2* is repressed in regions of high WntA signaling away from the wing margin (bottow). (**C-D’**) *WntA* mKOs show a distal shift of the blue dPF chevrons towards the submarginal band pattern (arrowhead). Wings with high mosaicism in the CSS (above the dotted line) show undulating chevrons in the dPf (D’). (**E-E’**) *fz2* mKOs phenocopy the distalization of dPf elements seen in *WntA* mKOs. The irregularity of mosaic clones suggests cell-autonomy (E’). (**F-G**) WntA-Fz2 signaling is active in presumptive CSS and marginal silver patterns of the posterior hindwing in *A. incarnata*. (**K-N**) Reversed deployment of WntA signaling around anterior silver spots in *A. incarnata* hindwings, explaining the expansion of silver spots in *WntA* KOs (N) that are anterior to the M_3_ vein (green). Scale bars = 1 mm.

### Inverted expression and patterning activity of WntA/Fz2 signaling across the M_2_ vein in *Agraulis*

The wing patterns of *A. incarnata* present an interesting conundrum on the ventral hindwing. As described above, WntA and Fz2 are both necessary for the induction of most CSS and peripheral silver patterns (Fig. 3F-J’). However, knock-outs of both genes result in an expansion and gain of silver spots anterior to the M_2_ vein (Fig. 1E) – for instance, the silver element situated in the M_1_-M_2_ region is normally split into two disjunct silver spots (dotted lines in Fig. 3M), while these spots fuse and extend basally in *WntA* and *fz2* KOs. Similarly, the M_2_-M_3_ region normally lacks silver elements (above the green line in Fig. 3M), but shows an ectopic silver pattern in these crispants. Why are there two opposed effects of WntA on silver spots? We previously speculated that WntA temporally shifted from activating to inhibitory role on silver spot formation, with the later corresponding to a burst of *WntA* signal observed in anterior hindwings shortly before pupation (Mazo-Vargas et al., 2017). Surprisingly, here we found that *WntA* expression is actually absent from the anterior silver spots, and is instead expressed in their brown contouring regions in pupal wings (Fig. 3K-N). WntA signaling is notably active in the domain that splits the M_1_-M_2_ into two spots, as observed by strong *WntA* and low *fz2* mRNA patterns, and decreased signaling results in silver spot extensions. In other words, while WntA/Fz2 generally activates silver and black outlines in *Agraulis* ventral surfaces, it can also locally shift to a complementary expression mode that induces the brown contours of similar patterns instead. These data establish WntA/Fz2 signaling as a versatile developmental tool for setting boundaries in a context-dependent fashion, with the capacity to induce elements that are variably perceived as “pattern” or “background” (Nijhout and Wray, 1988).

### Fz1 loss–of-function results in Planar Cell Polarity-like effects

Next, we assayed the other three Frizzled receptors for effects on development and wing patterning. First we generated *fz1* mKO knockouts across a large sample of *Junonia* embryos (N= 5,273), varying injection time, CRISPR duplex concentration, and testing two different sgRNA targets. We did not observe pattern effects in these butterflies, but instead, crispants showed disorganized arrays of scales on the wings as well as on antennae and abdomens (Figs. 4A-C’, S11). These effects are clonal (*i.e.* with visible boundaries between mutant and WT scale fields), and reminiscent of *Drosophila* PCP phenotypes (Adler, 2012; Lawrence and Casal, 2018): similarly to wing hairs in PCP mutant flies, *fz1*-deficient butterfly scales are globally misoriented, forming streaks of scales that deviate from the normal proximal-to-distal orientation. Mutant scales also appeared to take upright positions relative to the wing plane, indicative of a loss of asymmetry in the scale protrusion axis during early pupal development (Dinwiddie et al., 2014). Based on the fact that *fz1* is necessary for wing PCP in *Drosophila*, we propose butterfly scale phenotypes as mechanistically equivalent.

**Figure 4.**
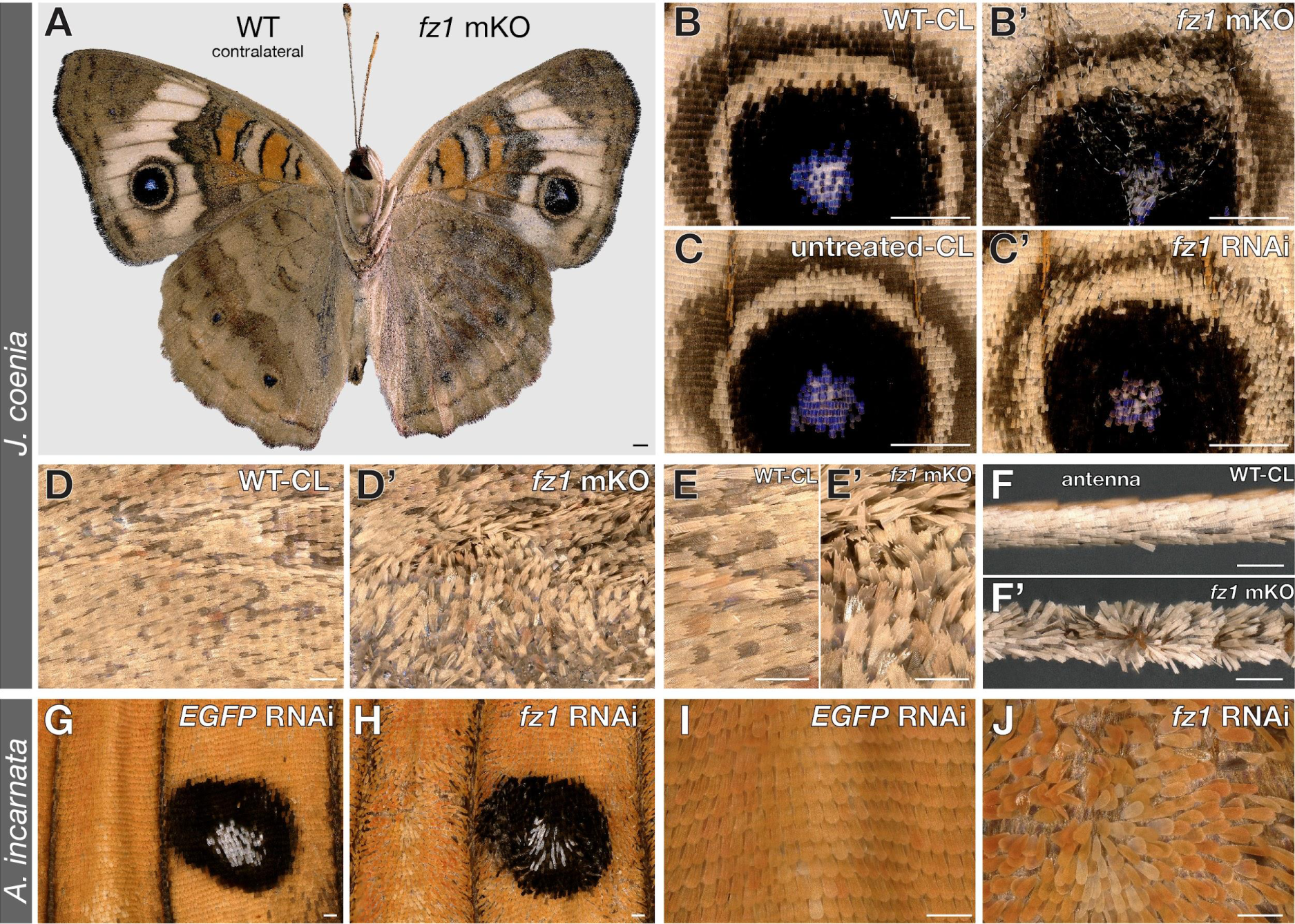
PCP-like scale orientation defects in *fz1* mKO and RNAi knockdowns. **(A)** *J. coenia fz1* crispant with mutant phenotypes limited to a single side (A: right, then magnified in B, D, E), and a contralateral wild-type (WT-CL) side (A: left, then magnified in B’, D’, E’, F’). **(B-B’)** PCP-like clones in the Cu_1_ eyespot from *fz1* mKO clones. **(C-C’)** Mild PCP-like phenotypes observed in electroporated RNAi experiments targeting *fz1*. **(D-E’)** Magnified view of an anterior forewing region comparing *fz1* mKO to WT-CL scales. WT-CL images were horizontally flipped to match the crispant wing direction. (**F-F’)** Scale orientation defects on antennae, as seen here in the asymmetric *fz1* crispant (A). **(G-J)** Strong PCP-like phenotypes observed after *fz1* siRNA electroporation in *A. incarnata* (N=8), compared to sham controls targeting an EGFP sequence (N=2). The Cu_1_-M_3_ ventral forewing CSS silverspot is shown in (G-H). Limb orientations: B-C’ and G-J, distal at the bottom; D-F’, distal on the right. Scale bars: A-C’ = 1 mm; D-J = 200 µm

Mosaic knock-outs of *fz1* in *Vanessa* also did not generate color pattern or scale defects among emerged adult butterflies (Table S1). However in contrast to *Junonia*, some pupae from this experiment showed wing growth defects and failed to form a cuticular seal with the abdomen, resulting in pupal death or failure of imago emergence due to desiccation (Fig. S12). Additionally, in the *Junonia* crispants that showed PCP phenotypes on one side only (Fig. S13), mutant forewings were shorter and rounder than their WT counterparts (Wilcoxon signed rank test, *p* < 0.05). This suggests that while *fz1* deficiencies can also affect wing growth in *Junonia*, pupal viability is less sensitive to perturbation in this species than in *Vanessa*.

To further verify its link to scale polarity, we used siRNA electroporation to generate knockdowns of *fz1* at the pupal stage (Fujiwara and Nishikawa, 2016; VanKuren et al., 2022), thus bypassing the wing growth phase from larval stages. Electroporated siRNA knockdowns consistently recapitulated PCP phenotypes in *Junonia* and *Agraulis* (Fig. 4C, G-J). In each butterfly, these effects were specific to the electroporated surface, and were not observed in sham-treated controls targeting a GFP sequence (Fig. S14). In *Junonia*, we also reproduced hindwing-to-forewing pattern homeoses by knocking-down the *Ultrabithorax* Hox gene (Komata et al., 2022; Tendolkar et al., 2021), as well as local removal of the CSS using *fz2* knockdowns (Fig. S15). These two positive controls both successfully reproduced expected effects and indicate that the PCP-like effect is not induced as an artifact of electroporation. Thus, the combination of CRISPR KOs and siRNA knockdowns indicate the conservation of Wnt-independent Fz1-PCP pathway that is required for the proper orientation of epithelial features, similarly to *Drosophila* (Ewen-Campen et al., 2020; Joyce et al., 2020).

### Fz3 and Fz4 are inhibitors of vein formation

Next, we found that mosaic knockouts of both *fz3* and *fz4* in *Vanessa*, *Junonia* and *Agraulis* generated similar effects on vein and color patterning (Figs. 5A-C, S16-20). Most notably, disruption of *fz3* and *fz4* result in ectopic veins across all wing surfaces, sometimes forming complete veins and sometimes forming short spurs or loops (Fig. 5D), showing that these two genes have non-redundant roles in the repression of vein formation. As the lacunae, which prefigure the veins, form during larval wing development, this vein inhibition function is likely to occur before metamorphosis (Banerjee and Monteiro, 2020; Nijhout, 1991; Reed and Gilbert, 2004; Reed et al., 2007). Consistently with this idea, *fz3* and *fz4* are expressed in the intervein regions of *Vanessa* larval wing discs (Fig. S1). In places where a supernumerary vein interacted with the eyespot field in *Vanessa* and *Junonia*, we observed alterations to the shape and size of eyespots, including the formation of additional eyespot foci (Figs. 5E, S16-19). This is consistent with mathematical models of eyespot formation where veins act as important signaling centers and domain boundaries (Connahs et al., 2019; Nijhout, 2017), and so is likely to be a secondary consequence of ectopic veins, rather than a direct interaction between Fz3/Fz4 and eyespot-inducing morphogens. Likewise, disruptions of CSS elements were accompanied by ectopic venation (*i.e.* Fig. 5A”), suggesting a collateral effect of venation on field boundary conditions rather than on the WntA morphogen itself.

**Figure 5.**
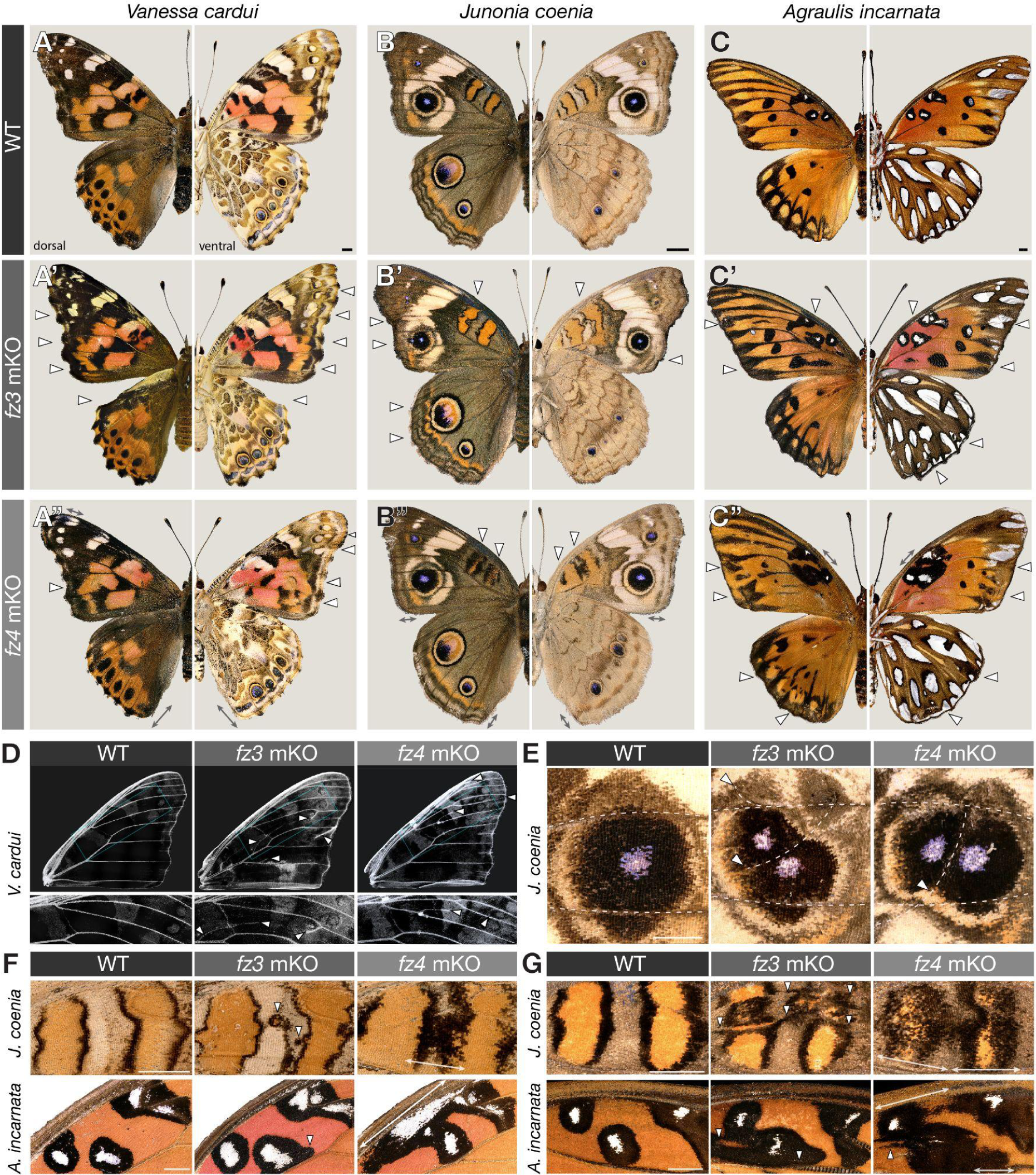
Effects of *fz3* and *fz4* mKOs on wing venation and color pattern formation. **(A-C’)** Dorsal (left) and ventral views) right of representative mosaic crispants for *fz3* and *fz4* in three species. Arrowheads: color pattern defects explained by ectopic venation; Double-arrows: cell non-autonomous color pattern defects observed without (A”, B”), or in conjunction with (C”) ectopic venation. **(D)** Both *fz3* and *fz4* mKOs induce ectopic veins (arrowheads), shown here in descaled *Vanessa* forewings. **(E)** Split eyespots observed in *fz3/fz4* crispants result from ectopic venation. (**F-G**) Ventral and dorsal Discalis pattern aberrations in *Junonia* and *Agraulis*. *fz3* crispants show irregular Discalis patterns with sharp boundaries, seemingly associated with ectopic venation (arrowheads). *fz4* mKO results in expansions of Discalis patterns with unsharp boundaries (double-arrows), suggesting cell non-autonomous effects on extracellular morphogens. Scale bars = 2 mm.

### Differential roles of Fz3 and Fz4 in color patterning and wing margin specification

We further examined possible color patterning functions of *fz3* and *fz4*, with their confounding effects on venation in mind. In *Junonia* and *Agraulis*, Discalis elements are bicolor symmetric systems and express Wnt ligand genes such as *wg* (Figs. S4I **and** S5B), *Wnt6* and *Wnt10* (Martin and Reed, 2014), and *WntA*, although the latter only contributes to the patterning of *Agraulis* D_1_ (Mazo-Vargas et al., 2017). Both *fz3* and *fz4* showed expansions of Discalis elements (Fig. 5F-G). These effects of *fz3* perturbation are again likely to derive from ectopic venation, as they are accompanied by visible epithelial thickening in these regions, or as they formed close to other veins that may have formed spurs. Interestingly, *fz4* mKOs had distinct effects that suggest a direct modulation of pattern-inducing morphogenetic signals, rather than a collateral effect of ectopic venation. In *fz4* crispants, D_1_ and D_2_ had expanded inner and outer elements on the ventral side, while on the dorsal side, only the outer black element was expanded (Fig. 5F-G). Unlike *fz3* phenotypes, these expansions appear broader and always include a graded, fuzzy effect at their borders, suggesting a possible cell non-autonomous effect on extracellular signaling. .

Observation of the wing peripheral regions of *Junonia* and *Vanessa* provide further insights (Fig. 6), as they juxtapose a sequence of multicolor patterns (dPF and MBS) with possible contributions of WntA and Wg/Wnt6/Wnt10 (Martin and Reed, 2014; Mazo-Vargas et al., 2017). Perturbation of *fz4* yields specific effects on MBS patterning, seen as a loss of the sub-marginal band in *Junonia* and an MBS distal shift or reduction in *Vanessa,* as well as a distalization of eyespots observed in *Vanessa* only (Fig. 6A-B, D-E, **and** G-H). In contrast, the color pattern arrangements of *fz3* crispants are similar to WT in complexity (*i.e.* similar sequences and sizes of color layers), but instead show distortions that indicate two kinds of tissue distortions: ectopic venation, and ectopic wing margins. These secondary wing margins are visible as distal outgrowths that feature the sequence of scale types normally found at the primary margin, including linear streaks of elongated margin scales (Fig. 6C, F **and** I). Together with the strong expression of *fz3* in the peripheral tissue of larval wing disks (Fig. S1F), these results suggest that *fz3* is necessary for proper margin specification.

**Figure 6.**
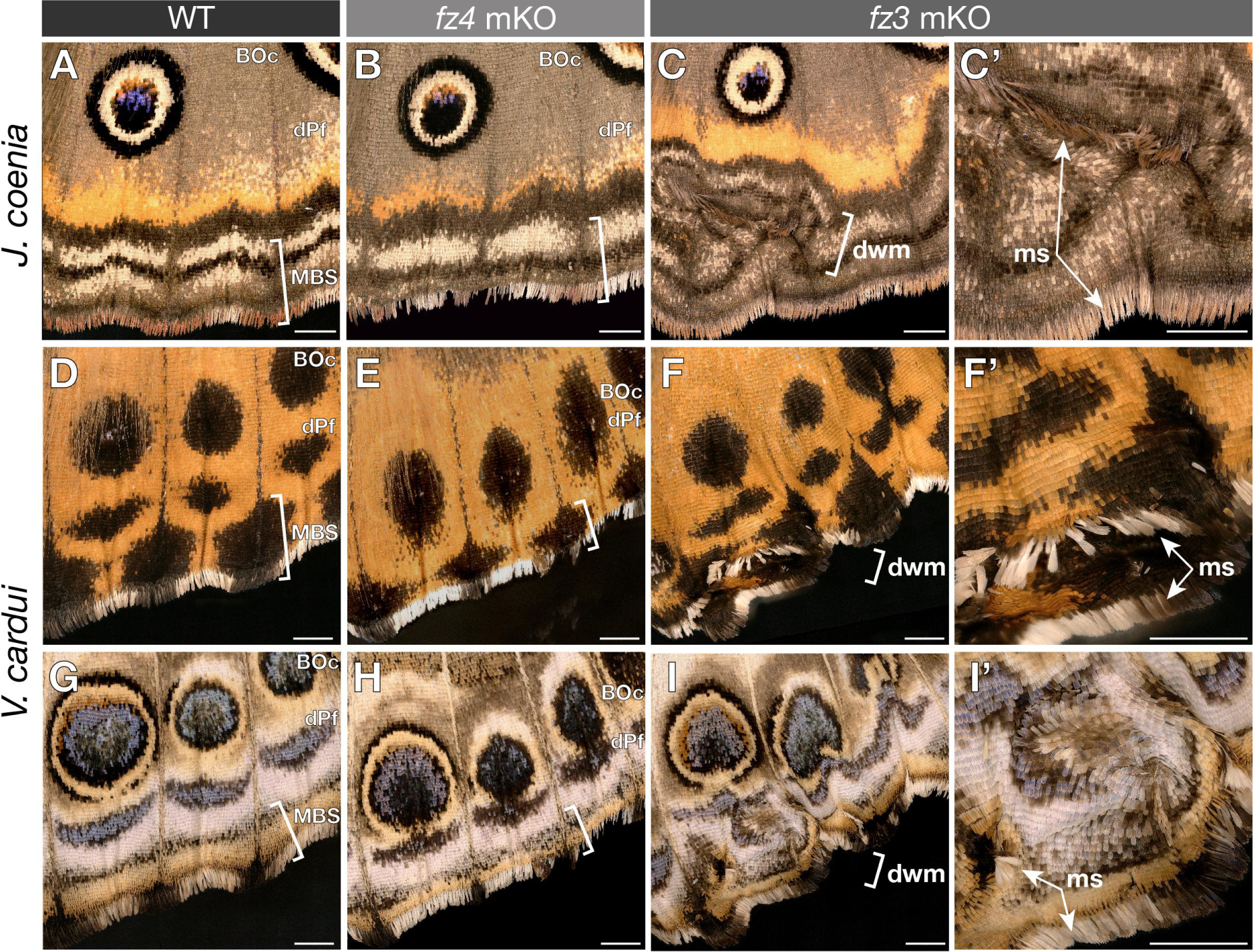
Peripheral effects of *fz4* and *fz3* mKOs on color patterns and the wing margin. **(A-C’)** Dorsal views of *J. coenia* crispants, with reduced MBS patterns in *fz4* mKOs, and localized distortions of the wing margin in *fz3* mKOs. In *fz3* mKOs, margin scales (ms, arrowheads) and concentric layering of color patterns and scale types around them indicate the presence of duplicated wing margins (dwm, magnified in C’, F’, I’), in addition to ectopic veinous tissue. **(D-I’)** Same observations in the dorsal (D-F’) and ventral sides (G-I’) of *V. cardui*, with the added finding that eyespots (Border Ocelli, BOc) are distally shifted and collapsing with the dPf chevron elements in *fz4* crispants of this species. Scale bars = 1 mm

## Discussion

### The WntA-Fz2 signaling axis instructs wing pattern formation

Here, we aimed to clarify the mode of reception of the WntA ligand, a knowledge gap explained by the evolutionary loss of *WntA* in both *Drosophila* and vertebrates. We showed that Fz2 is the sole Frizzled-family member necessary for WntA reception in butterflies. Importantly, both *WntA* and *fz2* knock-outs result in viable embryos, larvae and adults without deleterious effects on survival or morphology. It is interesting that these two genes are dispensable for normal embryonic development in spite of complex expression patterns (Hanly et al., 2021; Janssen et al., 2015). These results are in agreement with loss-of-function experiments in *Tribolium*, where knockdowns of *WntA* and *fz2* alone have no effect on embryogenesis (Beermann et al., 2011; Bolognesi et al., 2008), or in *Drosophila,* where null-homozygous *fz2* mutant adults are sterile but viable (Chen and Struhl, 1999; Cohen et al., 2002).

In addition, we gained insights on how *WntA*-dependent elements establish symmetric and complex arrays of color scales, such as the undulating brown patterns of the *Vanessa* CSS, or the *Agraulis* silver spots and their black contours (Figs. 2-3). We propose that WntA acts as a morphogen, inducing scale cell differentiation in a graded and concentration-dependent manner. First, WntA acts as a paracrine factor, as indicated by the cell non-autonomous nature of *WntA* crispant clones, with rounded boundaries that contrast with the expected cell-autonomous feature of receptor deficient clones in all species (Figs. 1G-J **and** 3). Second, we observe fine details in mRNA staining intensities that prefigure the differentiation of scales into multiple different types. For example, we see spatially intricate expression signals of *WntA* in the *Vanessa* hindwing ventral CSS: wild-type patterns display symmetric arrangements of beige, black, white, and brown scales, but in the WntA-deficient context, only one scale type remains (Fig. 3C-E). In *Agraulis WntA* KOs, the cell non-autonomous clonal effects we observed, such as the contraction of black-countoured silver spots into black spots, indicate a lower signaling level (Fig. 1G,I). This contraction is not observed in the corresponding patterns carrying cell-autonomous *fz2* mKO clones, where dislocated patterns feature both silver and black scales. These distinct phenotypes are consistent with receiving cells responding to specific thresholds in extracellular levels of WntA, or to duration of exposure, and rule-out sequential induction models involving intermediate signals (Iwata and Otaki, 2019). Together, the ability of WntA to act at a distance and to cause differentiation into multiple concentration-dependent cell types indicate that it is a morphogen.

We also observed localized depletion of *fz2* expression in WntA-dependent patterns, and a reduction of this effect upon *WntA* KO, showing that WntA/Fz2 high-level signaling triggers a negative feedback on *fz2* mRNA expression. We found that *WntA* expression is under positive feedback, a phenomenon that may locally amplify signal and notably refine expression levels that prefigure color outputs in the *Vanessa* CSS (Figs. 2A-H). This pathway-intrinsic dual feedback mirrors aspects of Wg/Fz2 regulation in the *Drosophila* wing disk, and is generally thought to mediate precise and robust developmental signaling (Lander, 2007; Mehta et al., 2021; Sagner and Briscoe, 2017). Overall, these results lay out an exciting avenue of research for studying the formation of discrete responses for graded input in the butterfly wing, an epithelial tissue that effectively “displays” differentiated cell states as scale colors, and for further understanding the dynamics of phenotypic evolution in such systems.

### Insights into Wnt pathway branching, a potential mechanism for wing modularity

In order to achieve the modular organization of pattern, butterfly wings are thought to deploy distinct cell signaling pathways and selector genes that control discrete sets of patterns in a regional-specific manner (Martin and Reed, 2014; McKenna et al., 2020). As multiple Wnt ligand genes are expressed in butterfly wing transcriptomes (Fenner et al., 2020; Hanly et al., 2019), it is important to elucidate how many Wnt pathways are active in color patterning, or in other words, if different branches of the Wnt transduction mechanisms can mediate separate cellular responses. Our current data offer a few insights. Strikingly, *WntA* and *fz2* KOs remove the brown CSS in *Junonia* but leave intact the overlapping, distinctively colored and *wg*-positive D_1_ and D_2_ (Fig. S5), that are thought to be induced by Wg/Wnt6/Wnt10 (Martin and Reed, 2014; Mazo-Vargas et al., 2017). It was also shown in the nymphalid *Bicyclus anynana* that Wg/Wnt canonical signaling is involved in the positioning of eyespots (Connahs et al., 2019; Özsu et al., 2017). We anticipate that *fz2* KOs alone would not impact eyespot development in *Bicyclus*, based on our findings that *fz2* loss-of-function did not affect eyespots in *Junonia* and *Vanessa*. These two sets of observations imply that different Wnt signals diverge in their effects across the wings and induce distinct responses, with WntA most noticeably inducing the CSS and dPF, and Wg (possibly with co-expressed Wnt6 and Wnt10) patterning Discalis, Border Ocelli, and MBS elements. However, the distinctiveness of these elements would be difficult to explain if color-patterning Wnt pathways converge on the same transduction mechanism. This leads us to the working hypothesis that the WntA/Fz2 pathway is non-canonical, potentially separating its signal from canonical Wg activity in different sets of patterns. Non-canonical Fz2 pathway branches have been identified in *Drosophila* and will deserve further examination in butterflies to test this possibility, including PTK7-family co-receptors (Linnemannstöns et al., 2014; Peradziryi et al., 2011), ROR-family co-receptors (Ripp et al., 2018; Shi et al., 2021), Fz2/Ca^2+^ signaling (Agrawal and Hasan, 2015), and the FNI pathway (Mathew et al., 2005; Restrepo et al., 2022).

### Evolutionary conservation of the Fz1-PCP pathway

Individual knock-outs of Fz1 and Fz2 failed to disrupt Wg-positive patterns, suggesting that Fz1 and Fz2 act redundantly to mediate canonical Wg functions in butterflies, similarly to *Drosophila* and *Tribolium* (Beermann et al., 2011; Cadigan et al., 1998; Ewen-Campen et al., 2020). It is plausible that this redundancy is not complete in certain tissue and species, potentially explaining wing growth defects of *Vanessa fz1* crispants (Fig. S7), as well as the larval cuticle defects observed in swallowtail *fz1* crispants (Li et al., 2015). Nonetheless, *fz1* knock-outs and knockdowns in several species showed that it is necessary for epithelial polarity with phenotypes displaying disorganized scale arrays akin to the wing trichome phenotypes characteristic of PCP in fruitflies (Adler, 2012). As in *Drosophila* (Ewen-Campen et al., 2020; Joyce et al., 2020), Fz1/PCP is independent from Wnt activity in butterflies, because knock-outs of the Wnt secretion factors Wls and Por do not yield scale polarity defects (Hanly et al., 2021). These data validate previous inferences that this pathway is functionally conserved across Bilateria and Cnidaria (Momose et al., 2012; Yang and Mlodzik, 2015), and prompt further studies of the Fz/PCP pathway in a wider variety of arthropods and tissue types. Future investigations of butterfly wing PCP in particular could look at the effect of *fz1* mutation on scale and socket precursor orientation, cytoskeleton distribution, and asymmetry of the Prickled PCP marker during the first phases of pupal wing development (Adler, 2012; Day et al., 2019; Dinwiddie et al., 2014).

### Fz3 and Fz4 are candidate Wnt decoy receptors

The expression patterns of *fz3* and *fz4* in arthropod embryos are diverse, but imply ancestral roles in nervous system and appendage patterning (Janssen et al., 2015). In *Drosophila,* Fz3 acts as a decoy receptor, *i.e.* as a competitive-antagonist that attenuates Wnt signaling and contributes to establishing the Wg gradient in wing discs (Sato et al., 1999; Schilling et al., 2014). Of note, *fz3* is lost from the genomes of *Tribolium* beetles and *Parasteatoda* spiders (Janssen et al., 2015). The *Drosophila* Fz4 binds the atypical WntD ligand to inhibit Toll signaling (Rahimi et al., 2016), but WntD is unique to the *Drosophila* lineage, and perturbation of *fz4* by RNAi knockdowns in *Tribolium* embryos did not identify a clear function (Beermann et al., 2011), meaning that Fz4 functions in other lineages remain elusive. In addition, both Fz3 and Fz4 have lost the intracellular Dsh-interacting domain (Fig. S1), supporting the hypothesis that they have non-canonical signaling functions. Here we showed that both *fz3* and *fz4* have key functions in the formation of normal venation in the developing butterfly wings. Clonal knockouts had sporadic, ectopic vein branching, implying that these genes are acting to repress vein development. No such role has been described in the literature for Fz3 or Fz4 in *D. melanogaster* (Ewen-Campen et al., 2020; López-Varea et al., 2021a). However, it is noteworthy that in *Drosophila* wing blades, activation of canonical Wnt signaling via misexpression of *wg* or of a constitutively active Arm induces ectopic veins (Brennan et al., 1999; Lunde et al., 2003; Zeng et al., 2008), and that knockdowns of the Wnt pathway genes *pegasus*, *Earthbound*, *pygopus* and *nemo* resulted in wing venation defects (López-Varea et al., 2021a; López-Varea et al., 2021b), overall suggesting that the Wnt pathway is indeed a modulator of vein formation in flies.

Similarly, we found that *fz3* is strongly expressed in the wing periphery and that its loss-of-function results in ectopic wing margin duplications (Fig. 6). Lepidopteran wing disks develop with their dorsal and ventral epithelia in apposition and form a wing margin, that separate pre-apoptotic peripheral tissue from the wing epithelium (Macdonald et al., 2010; Nijhout, 1991). The Wnt ligand genes *wg/Wnt6/Wnt10* as well as the transcription factor Cut mark the peripheral tissue, analogously to the Dorso-Ventral wing boundary (D-V) characterized in flies (Gieseler et al., 2001; Macdonald et al., 2010; Swarup and Verheyen, 2012). By analogy with the role of canonical Wnt signaling in specifying the D-V boundary in flies (Buceta et al., 2007; Swarup and Verheyen, 2012), it is plausible that the *fz3*-deficient secondary margins we observed in butterflies resulted from a Wnt signaling gain–of-function.

Together, these observations lead us to hypothesize that, as with *fz3* in *Drosophila*, both butterfly *fz3* and *fz4* are acting as decoy receptors that attenuate Wnt ligands by binding them without being structurally capable of transmitting a signal – in this model, the removal of one receptor would increase the amount of free Wnt ligand by reducing the ‘sink’ effect of the decoy, leading to higher Wnt pathway activity and thus the induction of ectopic veins. A similar mechanism may explain the effects of *fz4* knockout on Discalis and Marginal Band System pattern elements, which seemed to occur independently of venation defects. Specifically, the inner Discalis patterns (orange in *Junonia*, silver in *Agraulis*) expanded on ventral surfaces, while the outer black contours expanded in these same patterns on dorsal sides (Fig. 5F-G). In the peripheral patterns of both *Vanessa* and *Junonia*, we observed a distal shift of eyespots accompanied by a simplification of the MBS patterns. Importantly, both Discalis and peripheral pattern aberrations from *fz4* crispants in three species show rounded rather than jagged, irregular clones with sharp color breaks at boundaries (Figs. 5F-G **and** 6). We interpret these effects as cell non-autonomous, contrary to what would be expected from a classic receptor, but consistent with a Wnt decoy function occurring upstream of signal transduction. We thus hypothesize that Fz4 antagonizes Wnt in the extracellular space. It is worth noting that *fz4* mKOs resemble the effect of extracellular heparin injections in early pupae, a treatment that emulates Wnt gain-of-function, including Discalis element expansions (and injection timing influencing inner vs. outer pattern expansions in these), as well as MBS pattern fusion and expansions in *Junonia, Vanessa* and *Agraulis* (Martin and Reed, 2014; Serfas and Carroll, 2005; Sourakov, 2018; Sourakov and Shirai, 2020). Importantly, heparin likely enhances the effects of WntA in the CSS, but *fz4* mKOs only affected other heparin-sensitive patterns, implying that Fz4 is in fact unable to bind WntA. Instead, it is most likely that Fz4 is counteracting one or a combination of Wg, Wnt6 and Wnt10, which are all co-expressed in the Discalis elements and wing-peripheral region margins in larval wings (Martin and Reed, 2014). While expression and functional assays of Wnt-family members at the pupal stages will be needed to test this model, the current data suggest that sub-functionalized Fz3 and Fz4 antagonize distinct aspects of Wnt signaling in a context-dependent manner.

## Conclusion

The WntA/Fz2 signaling axis has heretofore remained unstudied due to the loss of WntA in classical model systems. But, it could well prove to be evolutionarily conserved, and it will be interesting to test if its role as a morphogen is involved in epithelial patterning elsewhere, or to identify other functions across Bilateria and Cnidaria (Hanly et al., 2021; Schenkelaars et al., 2015), such as its potential role in wound healing and regeneration (Li et al., 2017). The Fz1/PCP pathway and the Wnt-antagonist role of Fz3 both likely date back to prebilaterian ancestors (Momose and Houliston, 2007; Momose et al., 2012; Schenkelaars et al., 2015), while our sequence analysis suggests Fz4 only recently lost its Dsh-interacting domain in the course of insect evolution (Fig. S1, cyan box). Overall, functional analyses of Frizzled receptors, seen under the colorful features of butterfly wings, enrich our understanding of their diverse functions in mediating cell signaling and tissue patterning.

## Methods

### Butterflies

*Vanessa cardui* (Linnaeus, 1758) purchased from Carolina Biological Supplies, and *Junonia coenia* (Hübner, 1822), originating from the laboratory of Fred Nijhout (Duke University) were reared on artificial diet, in a growth chamber at 25°C, 40-60% relative humidity, and with a 14:10 h light:dark cycle, following previously described procedures (Martin et al., 2020; Tendolkar et al., 2021) with the following modification: to avoid the spread of viral disease, eggs are surface decontaminated for 2 min (*V. cardui*) or 4 min (*J. coenia*) with a Benzalkonium Chloride 5% solution, before being dried and left to hatch on double-tape placed under the cup lids, with no more than 40 eggs per container. In these conditions, the mean pupal developmental time is 160 h for *V. cardui*, and 190 h for *J. coenia*.

*Agraulis incarnata* (Riley, 1926) is the revised species name for subspecies of (formerly named) *Agraulis vanillae* found in the US (Núñez et al., 2022). *A. incarnata* adult butterflies were maintained in a closed enclosure situated in a rooftop greenhouse in Washington DC, with fluctuating temperatures 23-28°C and a periodic misting system for humidity. Gatorade 50% cups, *Lantana spp.*, and *Buddleia spp.* provided nectaring sources. Normal feeding and sexual behavior during the day required light supplementation with six Repti-Glo 10.0 Compact Fluorescent Desert Terrarium Lamp bulbs (Exo Terra). Eggs were collected on *Passiflora biflora* or *Passiflora incarnata* and washed 1 min with Benzalkonium Chloride 5%. Larvae were reared in an incubator at 28°C, 40-60% relative humidity, and with a 14:10 h light:dark cycle. This species is prone to viral disease when crowding occurs. To remediate this, larvae were reared on artificial diet, following the *V. cardui* rearing procedure with the following modifications. Smaller batches of embryos were hatched and reared during the initial stages (N < 8 individuals per cup), and larvae were moved to individual cup containers at the third larval instar. The Passionvine Butterfly artificial diet (Monarch Watch Shop), supplemented with dried *P. biflora* or *Passiflora incarnata* was used for larval feeding. Larvae required frequent movement to fresh diet cups, removal of excess silk in the fifth instar larvae, and addition of a horizontal toothpick section under the cup lid to induce pre-pupal silk pad spinning.

*Heliconius erato demophoon* butterflies (Ménétriés, 1855), and *Heliconius melpomene rosina* (Boisduval, 1870) were reared at the *Heliconius* insectaries in Gamboa (Panama) and at the LMU Munich using previously described methods (Concha et al., 2019; Rossi et al., 2020). *Danaus plexippus* (Linnaeus, 1758) butterflies were reared on milkweed in greenhouse cages and standard conditions at UC San Diego.

### CRISPR mosaic gene knock-outs

Egg microinjections were performed as previously described to deliver equimolar mixes of Cas9/sgRNA into butterfly syncytial embryos. In brief, sgRNAs were designed to target the transmembrane domain of the Frizzled proteins (sequences in Table S2), ordered as synthetic sgRNAs (Synthego), mixed in 1:2 mass ratio with recombinant Cas9-2xNLS (PNABio, or UC Berkeley Macrolabs-QB3), and microinjected in embryos at 1-6 hr AEL. The induction of frameshift mutations was verified in *V. cardui* and *D. plexippus* mosaic mutants for *fz2* (Table S3, **Fig. S_**).

### In situ hybridizations

Detection of mRNA expression was performed following a standard method (Martin and Reed, 2014), with minor modifications for early pupal wings. Pupal wings at 13-17% development tissues were dissected in cold 1X Phosphate Buffer Saline (PBS). A scalpel or straight Vannas Spring Scissors (8 cm / 2.5 mm tip) were used to cut the contour of forewing, which was then detached from the pupal case by first grabbing the base then pulling the wing carefully towards the distal edge, and forceps were used to remove a portion of the peripodial membranes while keeping for forewings and hindwings embedded at their edges. Forewings were placed back to the cuticle and transferred to fixative (1X PBS, 10 mM EGTA, 9.25% formaldehyde) for 30-40 min at room temperature, together with the entire pupal case with the hindwings attached. After fixing, wings were washed with cold PBT (1X PBS, 0.1% Tween20) twice while still embedded in their cuticle or pupal case, which were then removed with fine forceps. Subsequent PBT washes and the rest of the published protocol were carried out with two pupal wings per tube. For the final development of stains, wings were incubated in BM Purple (Roche Applied Science) for 10-15 h at room temperature. Riboprobes for *WntA* and *wg* were previously described (Martin and Reed, 2014). Riboprobes for the four *frizzled* genes were PCR amplified from *V. cardui* cDNA (Table S4), transcribed with a Roche T7 DIG RNA labeling kit, purified with Ambion MEGAClear columns, and stored at −80°C. Images were taken using a Nikon D5300 camera mounted to a Nikon SMZ800N trinocular dissecting microscope, equipped with a P-Plan Apo 1X/WF 0.105 NA 70 mm objective and a Nikon C-DS stand for diascopic mirror illumination.

### siRNA electroporations

Dicer-substrate siRNAs (DsiRNAs) were designed against nymphalid *fz1* and *fz2* coding sequences (Table S5) using both the online IDT siRNA designer tool and the siRNA iScore Designer web services (Ichihara et al., 2007), ordered at the 2 nmol or 10 nmol scales as Custom DsiRNAs with standard purification (IDT DNA Technologies), resuspended at 70-100 μM in 1X *Bombyx* injection buffer (pH 7.2, 0.5 mM NaH_2_PO_4_, 0.5 mM Na_2_HPO_4_, 5 mM KCl), and stored as frozen aliquots at −80°C until use. Electroporations were conducted within 30 min after the onset of pupation using a BTX ECM 830 electroporator with gold electrodes (Harvard Apparatus, Genetrodes bent gold tip, 5 mm). Electroporation procedures followed a previously described procedure (Fujiwara and Nishikawa, 2016). In brief, pupal forewings are lifted over a thin pad of 1% agarose prepared in 10X PBS, injected in their ventral surface with 1-4 µL of annealed siRNA mix (distal to the discal vein), and covered with a 20-30 µL droplet of 1X PBS. The positive electrode is placed in contact with the droplet, taking great care to not touch the wing epithelium (see below), while the negative electrode is in contact with the agarose pad on the dorsal side of the wing, before electroporation with 5 square pulses of 280 ms at 9-15 V, separated by 100 ms intervals. The PBS droplet is removed before reinsertion of the forewing into its pupal case. Pupae are sprayed with water every two days before emergence to avoid desiccation. In order to control for potential artifacts of the procedure, sham controls using GFP siRNA injections and injection dye were run. GFP siRNAs were injected into forewings or hindwings and electroporated with 5 square pulses of 280 ms at 8-15 V, separated by 100 ms intervals. Similarly, Phenol Red (2-4 μl) was injected at a concentration of 0.05% to monitor injection spread across the forewing. In the red dye injection experiments, to test for potential effects of excessive current, the positive electrode was directly placed in contact with the wing tissue within a PBS droplet, before electroporation using 6V to 9V at 280 ms, 5 pulses with 100 ms interval each.

### Wing vein imaging

Wings were de-scaled by washing in 90% EtOH for approximately 3 minutes, then switching to 10% Clorox bleach for 3 minutes, and repeating until the scales and pigments were removed from the wings. Washed wings were then imaged on a Keyence VHX-5000 microscope mounted with a VH-Z00T lens.

## Authors Contributions

Conceptualization, Visualisation and Writing, JJH, LSL, AM. Investigation, JJH, LSL, AM-V, TSR-M, LL, AT, CRD, NL, EAE, OBWHC, ND’S, JJH-P, CM, JA, MR, MWP, AM. Supervision, JJH, WOM, MWP, AM. Funding Acquisition, JJH, WOM, AM.

## Acknowledgements

We thank Rachel Canalichio and the staff of the Wilbur V. Harlan Greenhouse at the GWU for butterfly rearing and host plant resources; Riccardo Papa, Bob Reed, Brian Counterman, and Donya Shodja for stimulating discussions and comments on the manuscript; Chip Taylor and Ann Ryan for providing Passionvine Butterfly artificial diet, and Richard Merrill for facilitating *H. melpomene* experiments. This work was funded by the NSF grants IOS-1656553 and IOS-2110534 to AM, the Smithsonian Biodiversity Genomics Postdoctoral Fellowship to JJH, Harlan Summer Fellowships to LSL, AT, EAE and OBWHC, a Luther Rice Fellowship to JJH-P, an NSF PRFB fellowship to AMV, and the DFG grant ME4845/1-1 to Richard Merrill.

## Supplementary information

**Figure S1.**
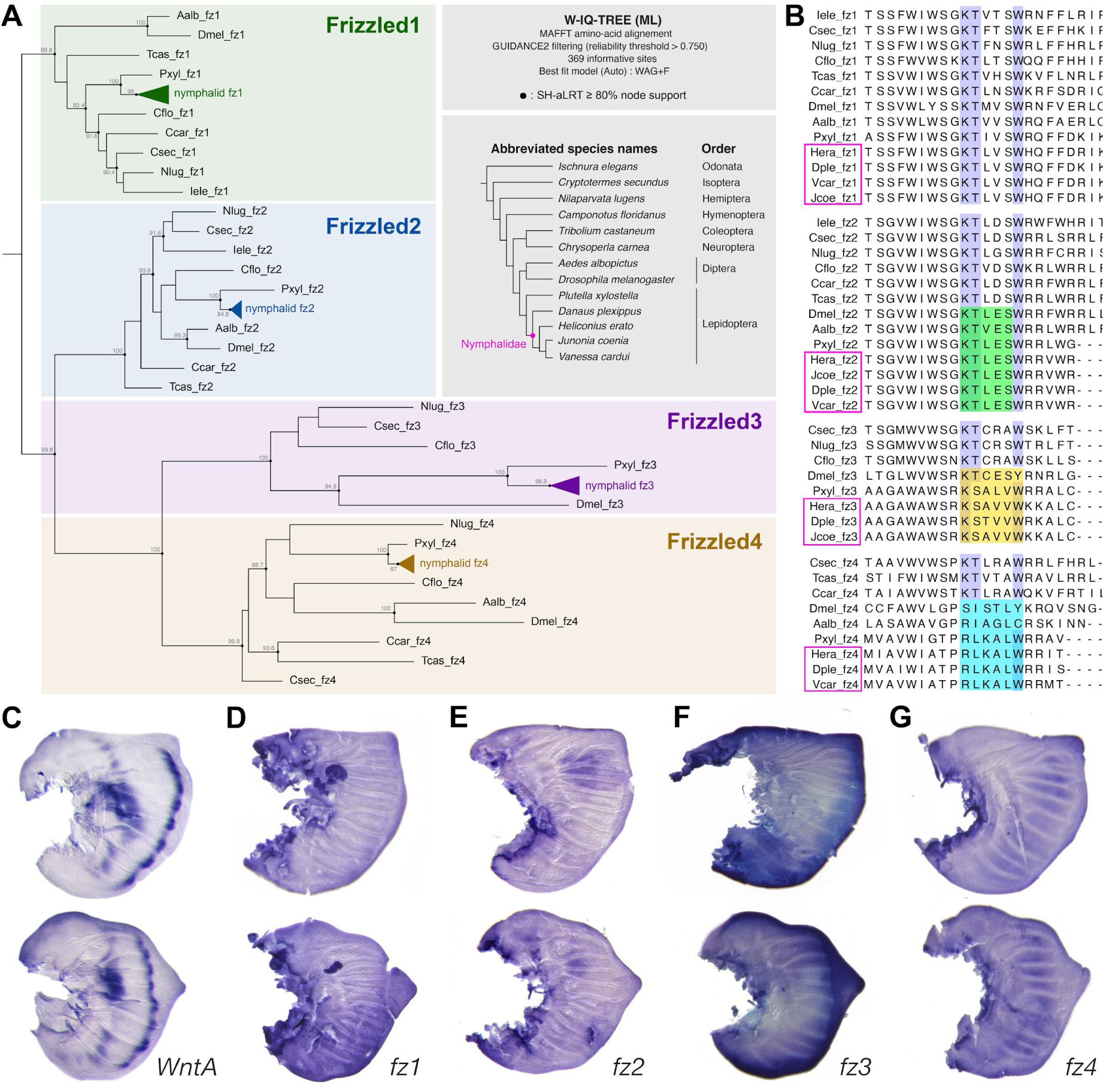
Four Frizzled-family receptors in Nymphalidae. (A) Phylogenetic reconstruction of insect Frizzled family receptors (source sequences and accession numbers in Supplementary File 1). Nymphalid butterflies encode 4 *frizzled* genes belonging to pre-Bilaterian orthology groups (Schenkelaars *et al*. 2015). **(B)** Alignment of insect Frizzled genes spanning the end of the TM7 domain (positions 1-7) and the beginning of the C-terminal cytoplasmic region (positions 8-23). Magenta: nymphalid butterfly sequences. Blue: conserved KTxxxW motif necessary for interaction with Dsh. Green: both lepidopteran and dipteran genomes show a conserved KTLES motif, needed for the Dsh-independent FNI transduction of Fz2 in *Drosophila* (Matthew *et al*. 2005). Yellow: Fz3 shows degeneracy of the KTxxxW motif in Lepidoptera and Diptera. Cyan: the KTxxxW motif is absent from lepidopteran and dipteran copies of Fz4. **(C-G)** *In situ* hybridizations of mRNA probes in *V. cardui* 5th instar larval imaginal disks for forewings (top) and hindwings (bottom). **(C)** *WntA* shows strong expression in the presumptive central symmetry system and marginal band system of larval wing discs. **(D)** *fz1* shows low ubiquitous expression throughout the imaginal disk. **(E)** *fz2* shows low overall expression with increased expression anterior of the M_3_ vein. **(F)** *fz3* shows intense staining in the *wg-Wnt6-Wnt10* expressing peripheral tissue, with weaker expression visible in the wing epithelium. **(G)** *fz4* shows interveinous expression.

**Figure S2.**
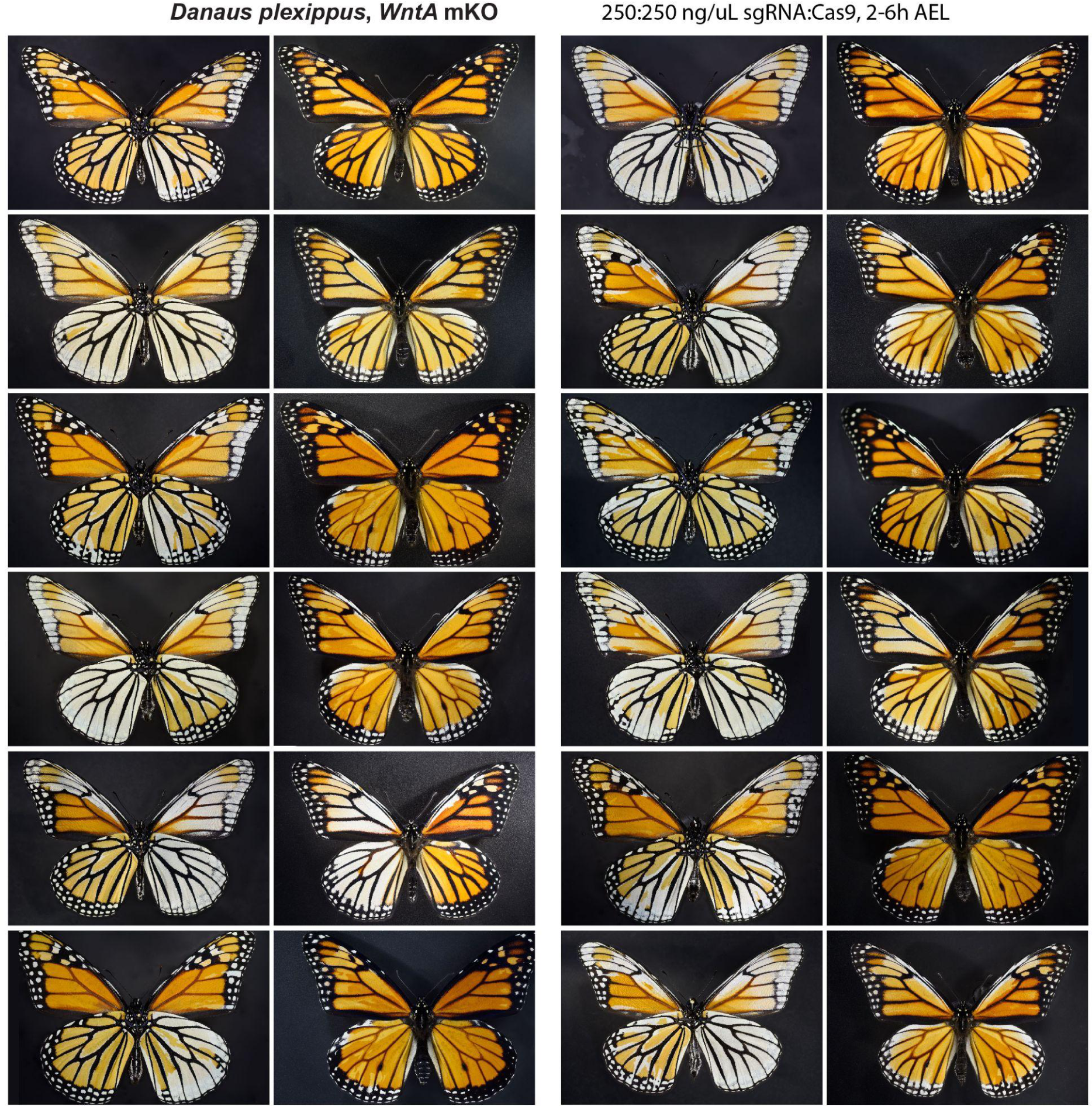
Representative *D. plexippus WntA* crispant phenotypes. Ventral (left image) juxtaposed to dorsal (right image) sides for each individual.

**Figure S3.**
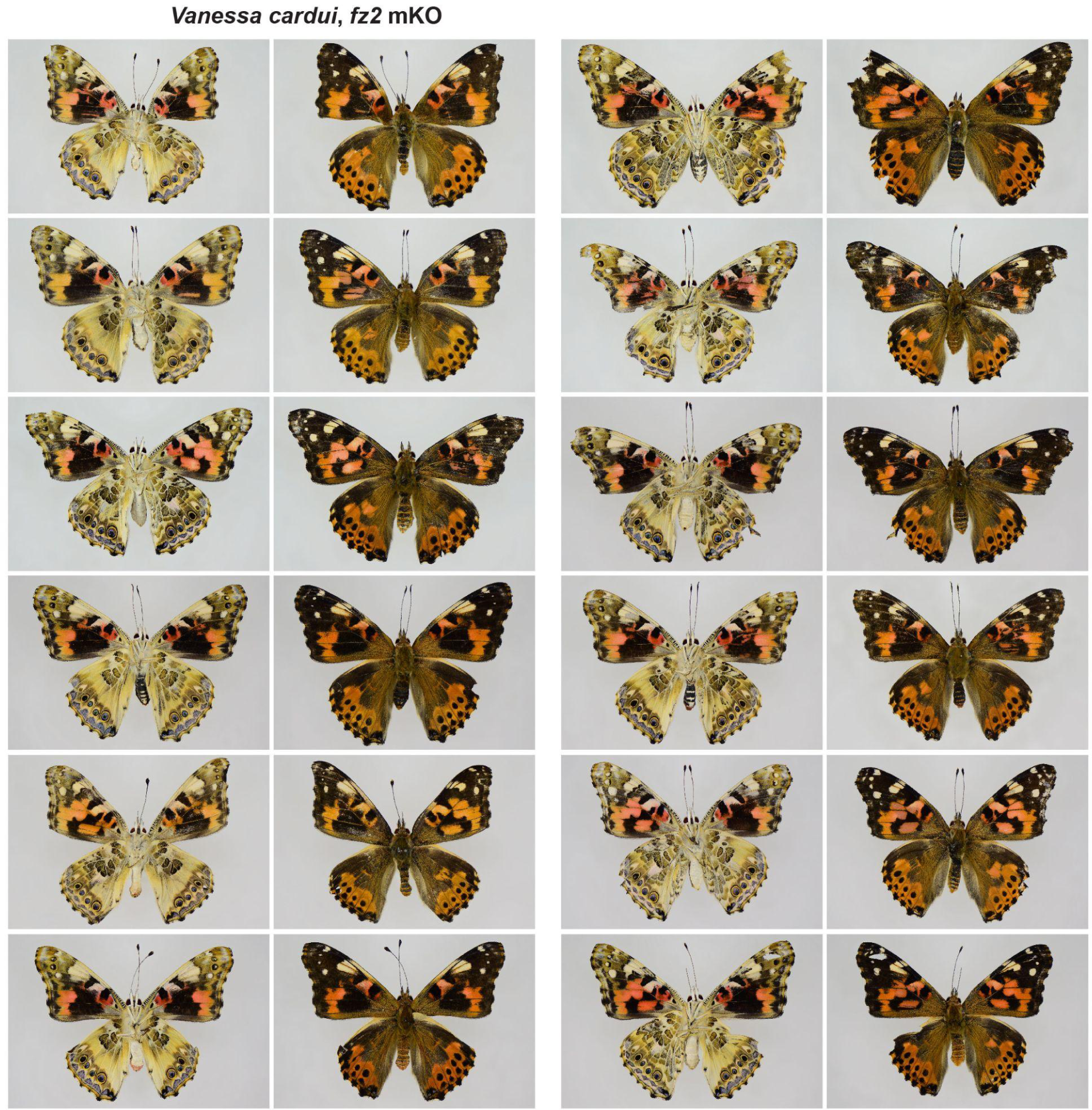
Representative *V. cardui fz2* crispant phenotypes. Ventral (left image) juxtaposed to dorsal (right image) sides for each individual.

**Figure S4.**
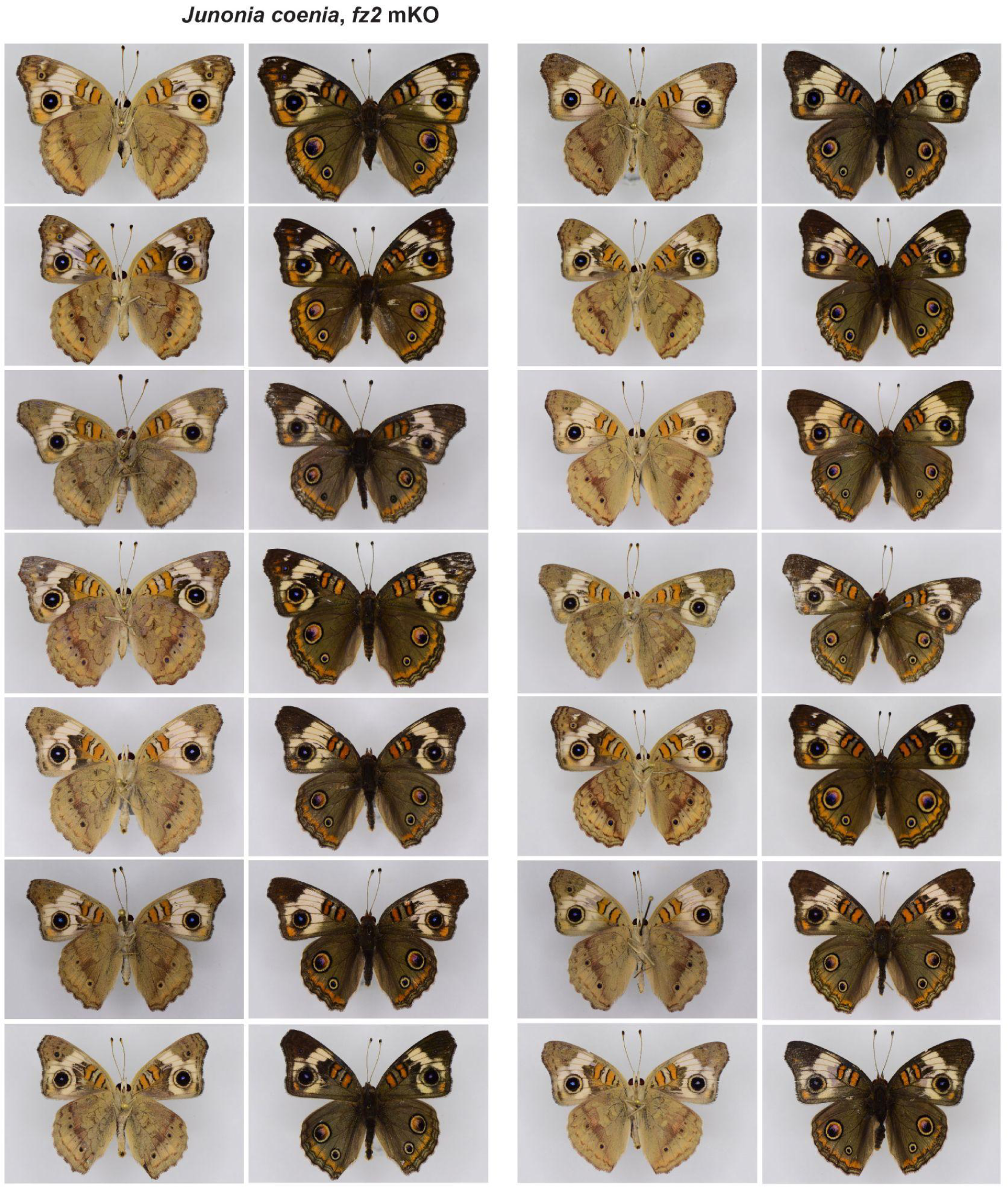
Representative *J. coenia fz2* crispant phenotypes. Ventral (left image) juxtaposed to dorsal (right image) sides for each individual

**Figure S5.**
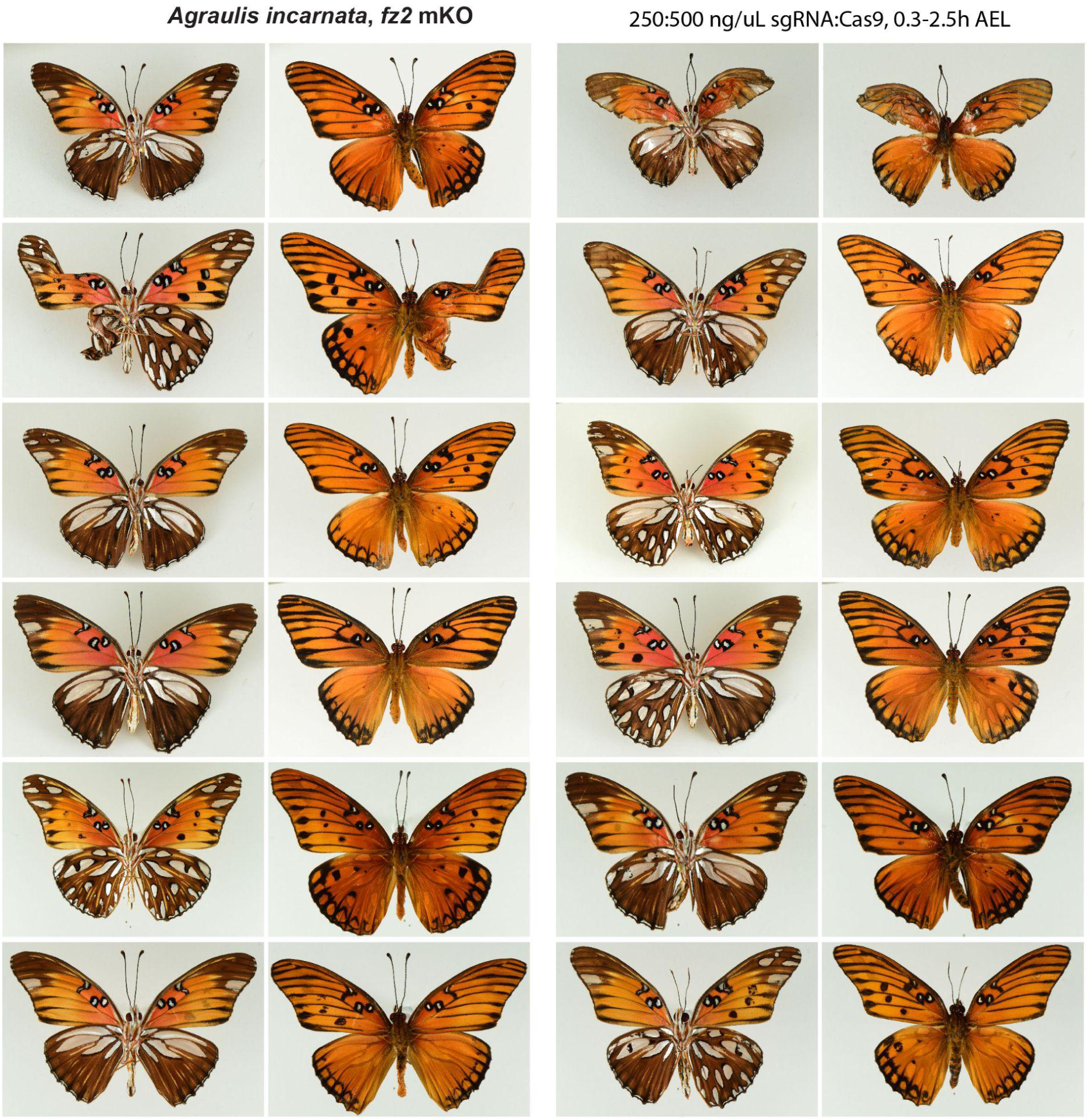
Representative *A. incarnata fz2* crispant phenotypes. Ventral (left image) juxtaposed to dorsal (right image) sides for each individual.

**Figure S6.**
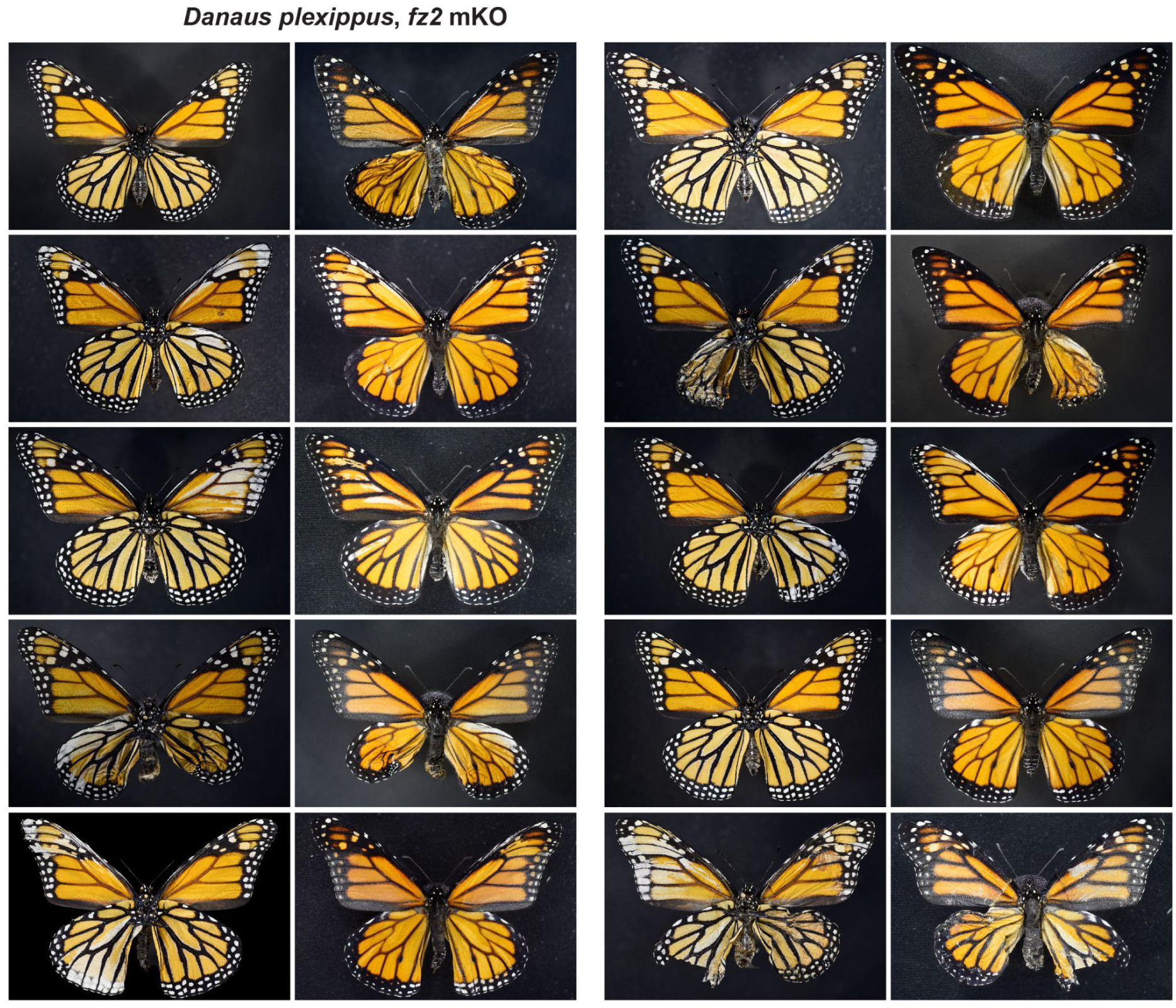
Representative *D. plexippus fz2* crispant phenotypes. Ventral (left image) juxtaposed to dorsal (right image) sides for each individual.

**Figure S7.**
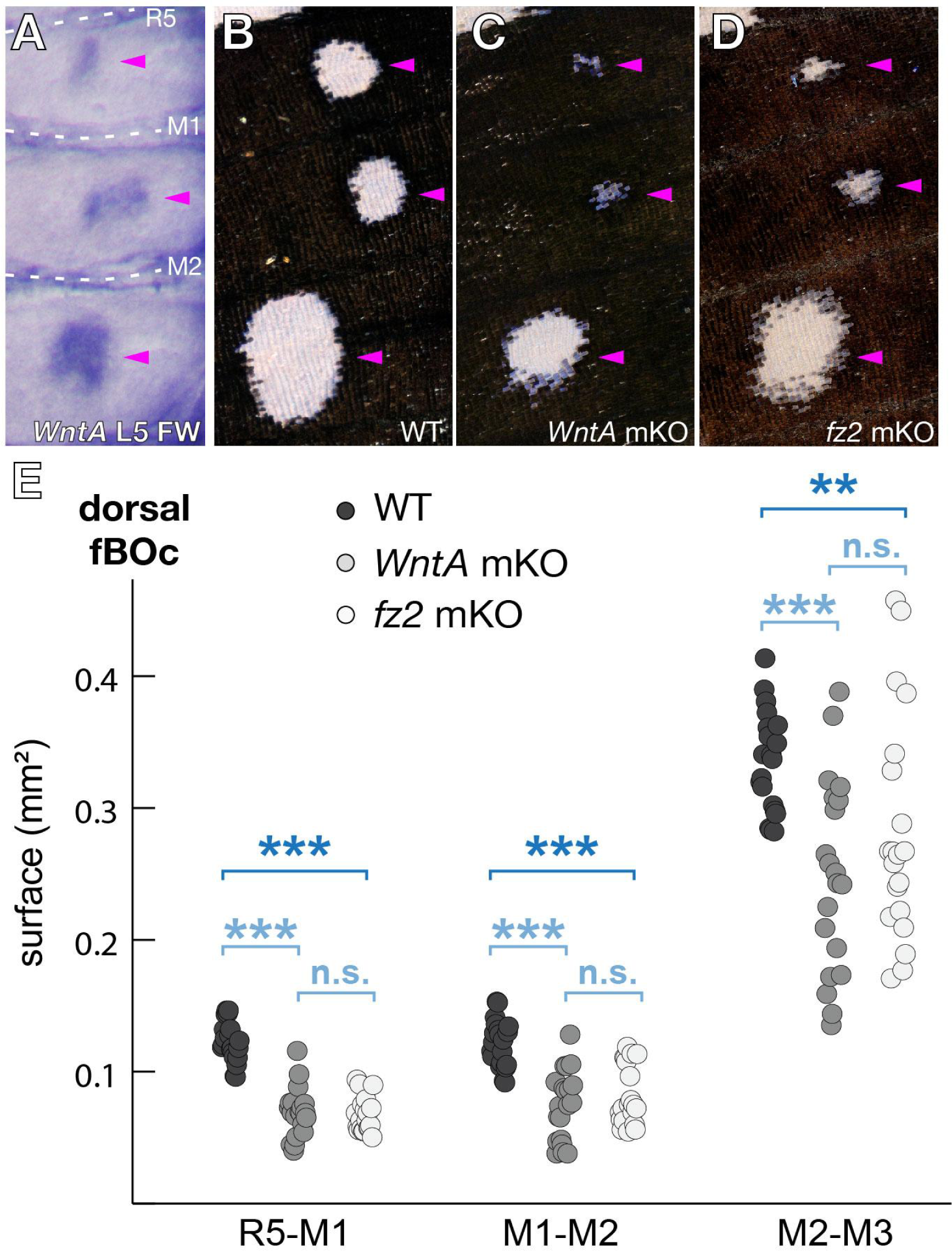
*fz2* mKOs phenocopy *WntA*-deficient reductions in the forewing Border Ocelli (fBOC) of *V. cardui.* (A) *In situ* hybridization showing *WntA* expression in the presumptive fBOc of *V. cardui* (arrowheads), situated between the R_5_-M_3_ veins, in late fifth instar wing disks. This fBOc expression persists in pupal stages (Fig. 2), and has not been described in nymphalids other than *Vanessa* to date. **(B-D)** Reduction of fBOc compared to WT in both *WntA* and *fz2* crispants, here in dorsal views. **(E)** Surface measurements of dorsal fBOc reveal significant reduction in both *WntA* and *fz2* mutant contexts (Mann-Whitney U-tests, **: *p* <0.01; ***: *p* <0.001; n.s.: non-significant). Figure adapted from (Mazo-Vargas et al., 2017) with *fz2* data added.

**Figure S8.**
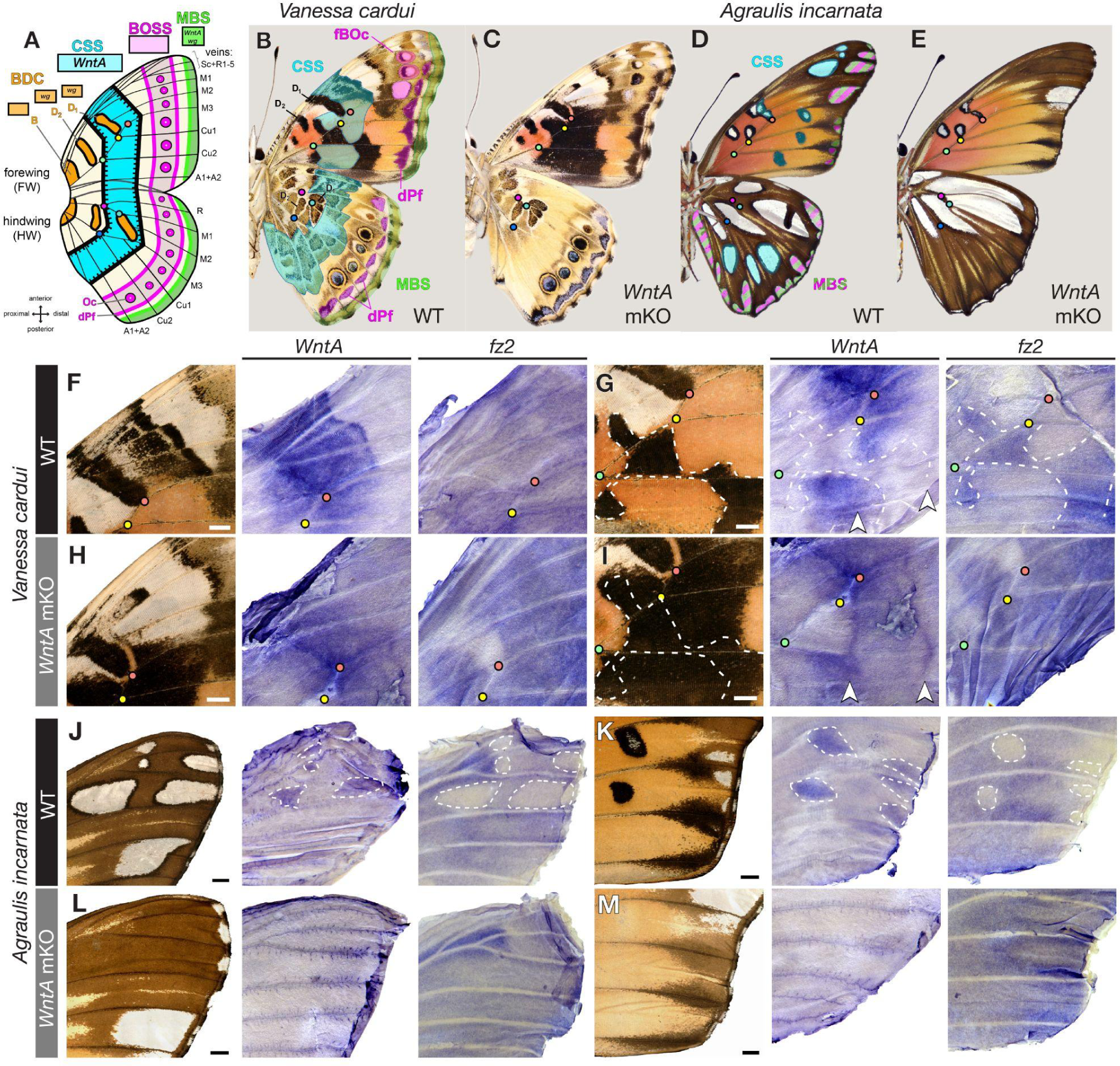
Expression of *WntA* and *fz2* in pupal wings is under the control of positive and negative feedback. **(A)** The Nymphalid groundplan (NGP) consists of four symmetry systems: Baso-Discal Complex (BDC, orange), Central Symmetry System (CSS, cyan), Border Ocelli Symmetry System (BoSS, magenta), and Marginal Band System (MBS, green). Color dots mark vein intersection landmarks (red: crossvein-M_3_; yellow: M_3_-Cu_1_; green: Cu_1_-Cu_2_). **(B)** Derivation of the NGP in ventral *V. cardui*. fBOc: forewing Border Ocelli; dPf: distal Parafocal elements. **(C)** *WntA* loss-of-function impacts the *Vanessa* CSS, fBOc, dPF, and MBS patterns. **(D)** Derivation of the NGP in ventral *A. incarnata*. The CSS is dislocated, and marginal patterns may include partial homology with dPF elements (hashing). **(E)** *WntA* loss-of-function impacts the *Agraulis* CSS and marginal patterns, and expands the anterior hindwing silver spots. **(F-I)** ISH for *WntA* and *fz2* mRNA across WT and *WntA* crispant wings at 13-17% pupal development (N = 3-4 replicates per experiment). In *WntA* mKO, *WntA* is modified (*e.g.* arrowheads), and *fz2* is de-repressed and becomes ubiquitous. (**J, L**) *Agraulis* forewing anterior CSS. (**K, M**) *Agraulis* posterior forewing and margin featuring the two CSS silver spots that flank the Cu_1_ vein. Contralateral *WntA-fz2* stains were obtained for F-I and K. Scale bars = 1 mm.

**Figure S9.**
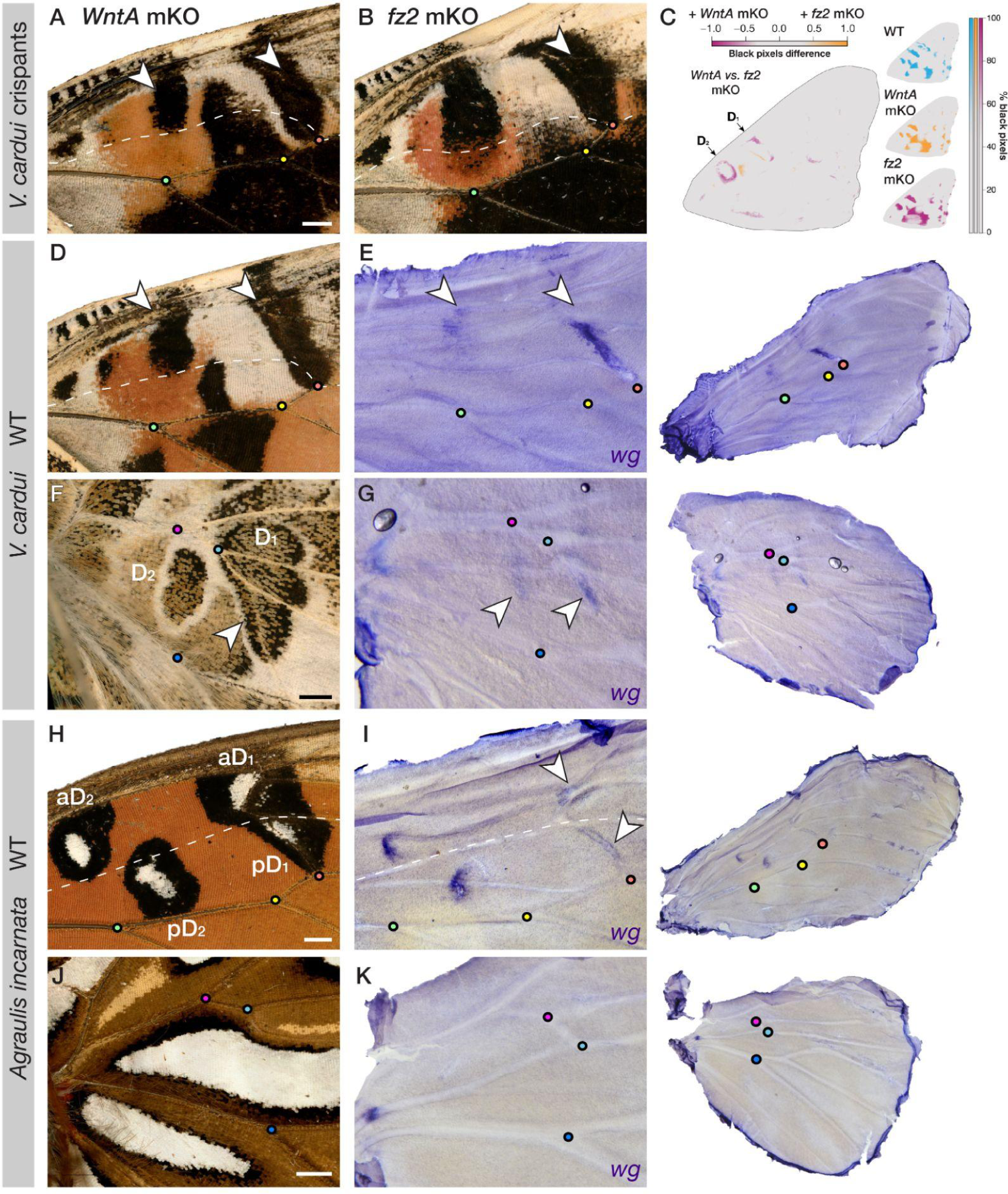
Size regulation of Discalis elements may involve Wg/Fz2 signaling in ventral *V. cardui* and *A. incarnata* forewings. **(A)** The D_1_ and anterior D_2_ Discalis (aD_2_) elements are unaffected by *WntA* mKOs. **(B)** D_1_ and aD_2_ show an expansion of black in *fz2* KOs. **(C)** *R/patternize* heatmap comparing *WntA* vs. *fz2* mKO forewings. Black scale expansions were consistently observed in D_1_ and D_2_ across all the *fz2* mKO samples (arrows). The upper row features black pixel distributions across WT (N = 20 images), *WntA* mKO (N= 20), and *fz2* mKO (N= 40). **(D)** Pupal forewing expression of *wg* marks presumptive aD_2_ and D_1_ patterns, suggesting exaggerated *fz2* phenotypes may be due to a role of Wg/Fz2 signaling in inhibiting their size. **(E)** Pupal hindwing expression of *wg* marks the Basalis (B), D_2_, and D_1_ elements. These three elements remain after *WntA* and *fz2* loss-of-function. **(F)** Pupal forewing of *A. incarnata* is marked by *wg* in the presumptive D_2_ and D_1_ patterns similar to in *V. cardui.* **(G)** Pupal hindwing expression of *wg* is only in the presumptive B element. **(H-K)** In *A. incarnata* 15-17% pupal wings, *wg* is most prominent in the forewing D_2_ (split across the anterior and posterior compartments, here marked with the M_2_ vein as dotted line), weakly expressed in the forewing D_1_, and absent from hindwings where Discalis elements are missing. Scale bars = 1 mm

**Figure S10.**
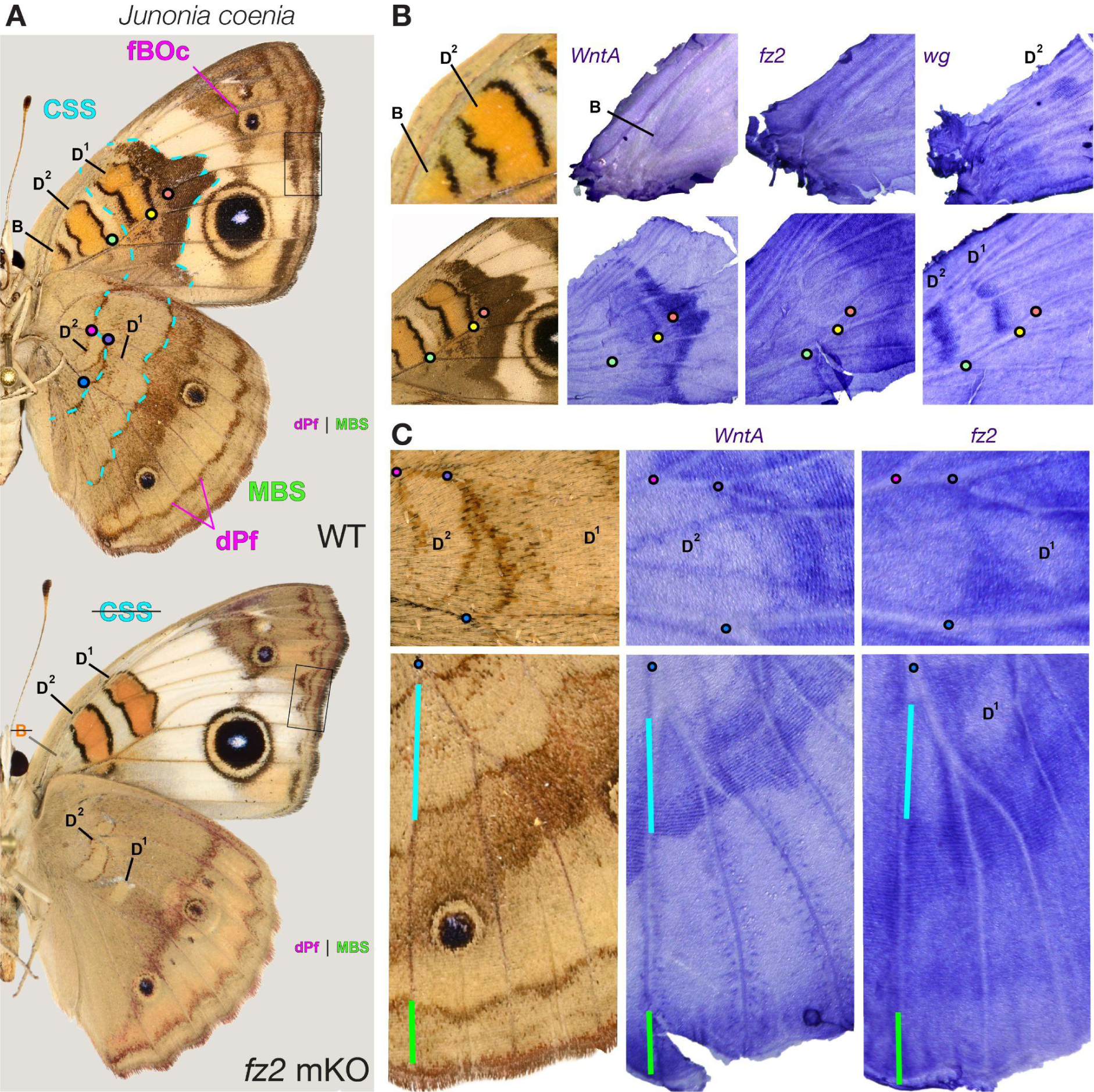
Conserved functions of *WntA* and *fz2* in a third nymphalid butterfly. **(A)** *J. coenia* wild type with annotated NGP elements and *fz2* mKO mutant. *fz2* crispants show a complete loss of the Basalis (B) and CSS elements, as well as a distalization and reshaping of the dPF elements, most visible in the forewings (box). **(B)** *In situ* hybridizations of *WntA*, *fz2* and *wg* in the proximal regions of the *J. coenia* WT forewing at 17% pupal development (n = 5 replicates). *WntA* induces the B and CSS elements, with visible *fz2* depletion at this stage, consistently with a negative feedback of WntA/Fz2 signaling on *fz2*. wg is strongly expressed in D_1_ and D_2_. **(C)** Expression of WntA and depletion of fz2 in the hindwing CSS (cyan) and MBS region (green) is consistent with a role of the WntA/fz2 signaling on inducing the CSS and positioning peripheral patterns.

**Figure S11.**
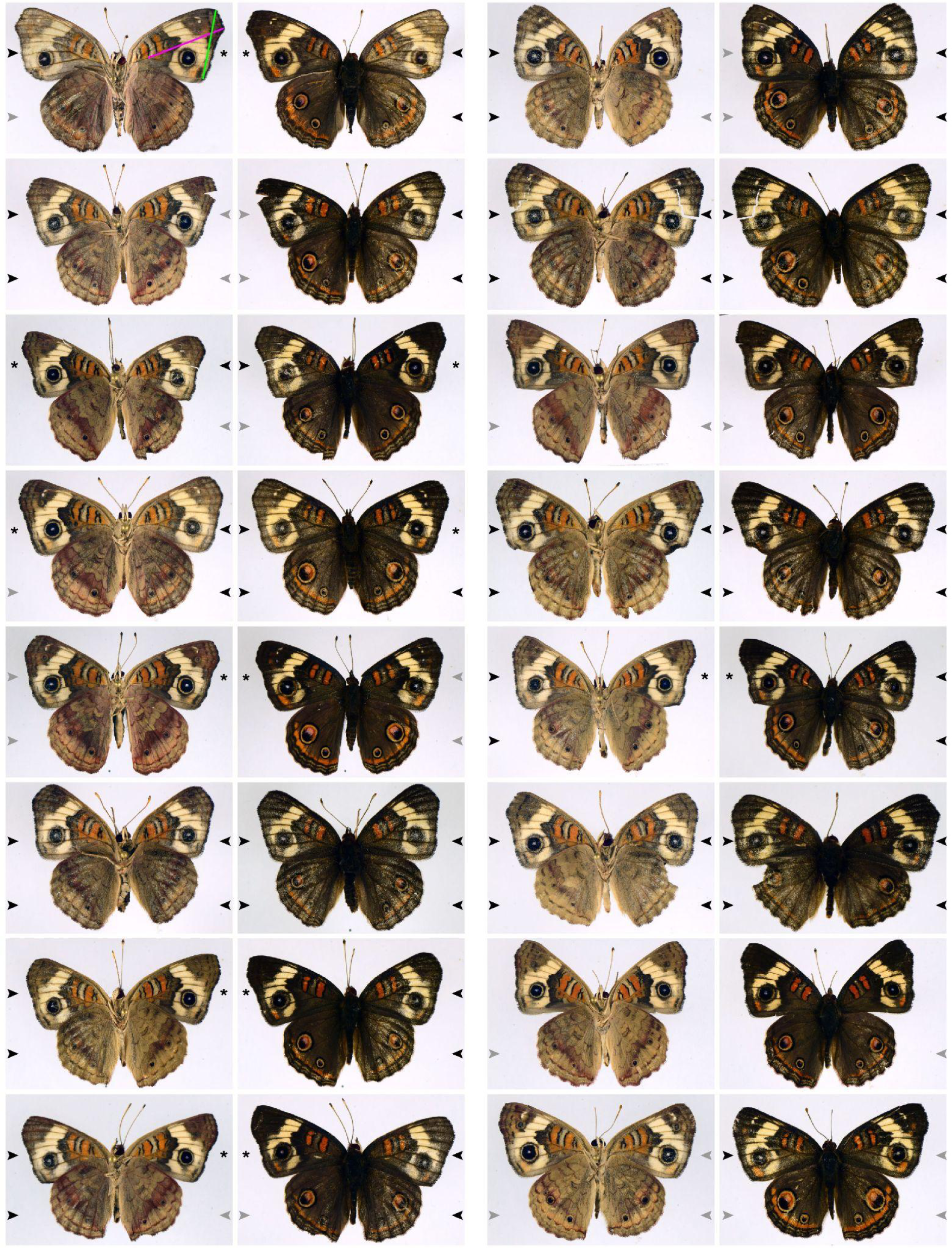
Mosaic *fz1* crispants show wings with disorganized scale arrays, short forewings, and no pattern defects in *J. coenia*. PCP phenotypes are visible in these whole-specimen views as zones of apparent wing wear. Left: ventral side; right: dorsal side. Black arrowheads: extensive PCP phenotypes; gray arrowheads: wing surface with small PCP clones; no arrowhead: surface with WT phenotype. Asterisks denote forewings that are WT on ventral and dorsal sides, and contralateral to a PCP-mutant forewing. Wing length (magenta) and width (green) between vein landmarks were measured in these PCP-asymmetric forewing pairs (N=7). Mutant wings are significantly shorter than WT in length (Wilcoxon signed rank test, *p* = 0.039) but not in width (*p* = 1).

**Figure S12.**
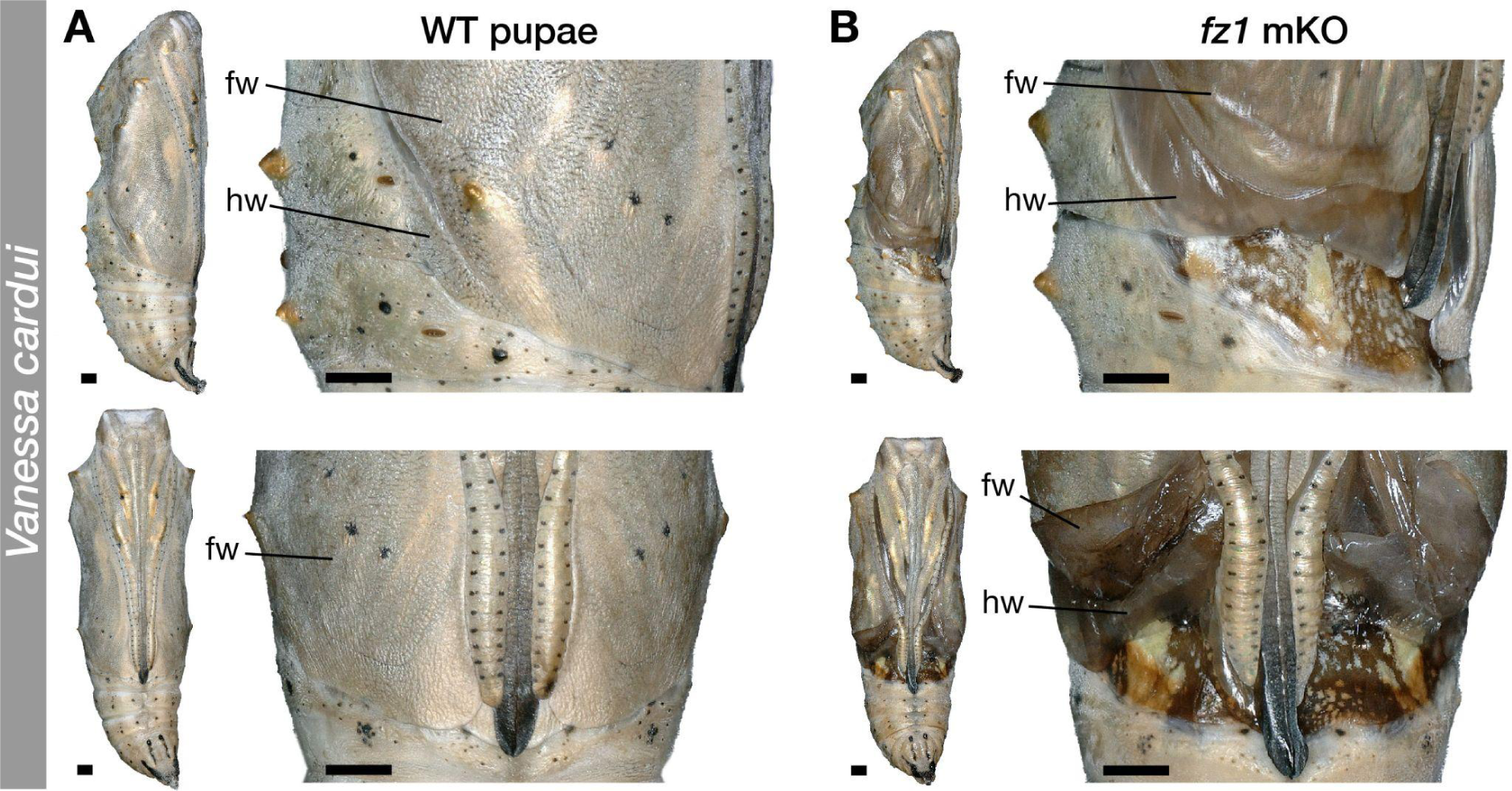
Pupal wing defects with incomplete wing growth and cuticularization in *fz1* crispants of *V. cardui*. **(A)** WT pupa in lateral (top) and ventral (bottom) views, with insets showing the distal portion of the pupal forewing (fw) and hindwing (hw) suturing with thoracic and abdominal cuticle. (**B**) Two examples of pupal phenotypes observed in *V. cardui fz1* crispants (N=13), with incomplete wing growth and distal sealing. Antennae, proboscis and legs, which bundle around the ventral midline, appeared unaffected. Scale bars = 1 mm

**Figure S13.**
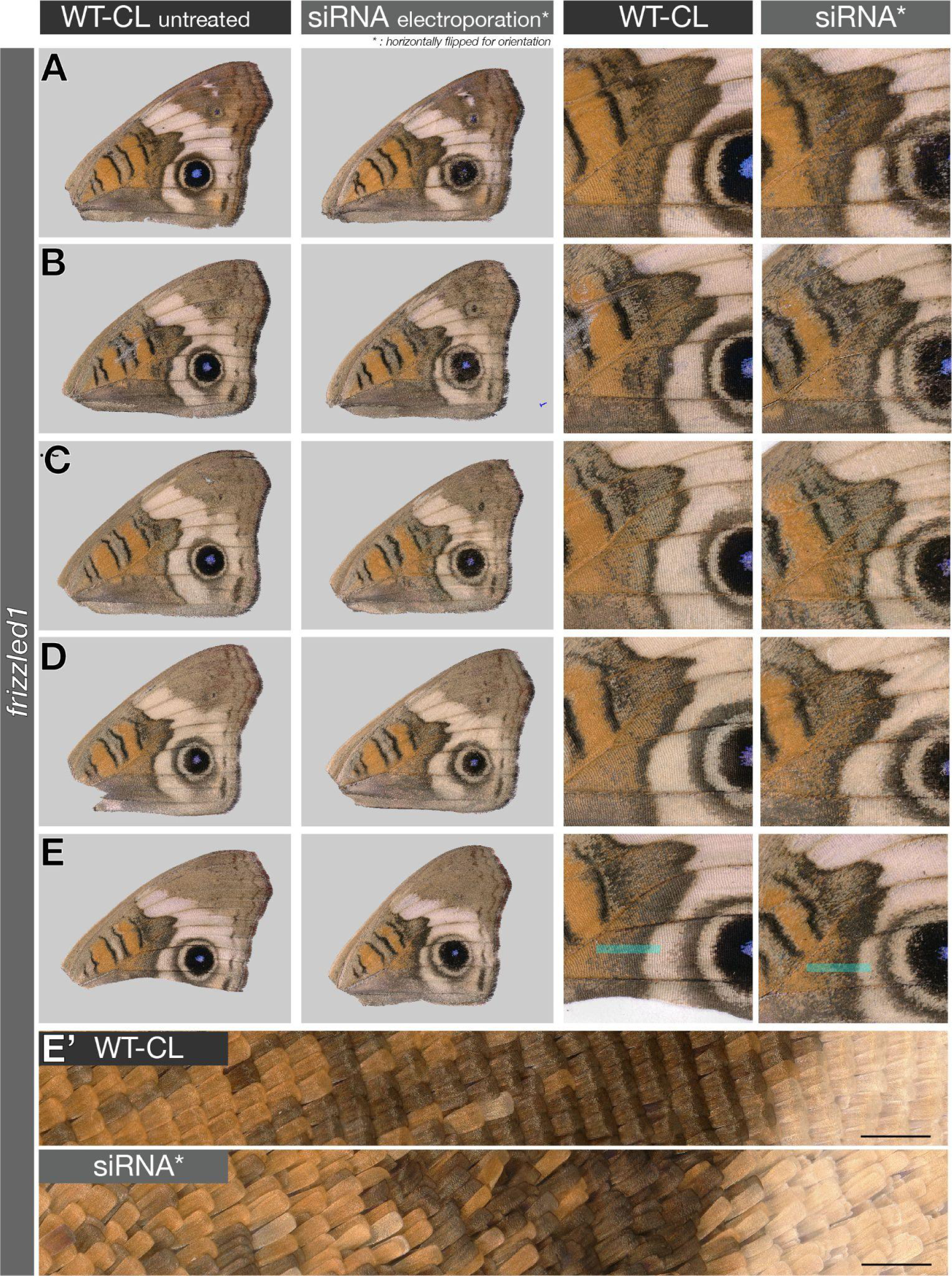
Pupal RNAi electroporation knockdowns of *fz1* in *Junonia.* Ventral views of single specimens electroporated with *fz1* DsiRNA on the ventral right forewing, shown here next to their contralateral wild-type control (WT-CL). PCP-like phenotypes are visible as fields of straightened scales (lacking normal curvature) and lacking regular organization under high magnification in all five replicates (as shown in E’, bottom panel) and were not observed in controls. PCP-like phenotypes were pronounced in *Agraulis* knockdown experiments (Fig. 4G-J). Scale bars: E’ = 200µm.

**Figure S14.**
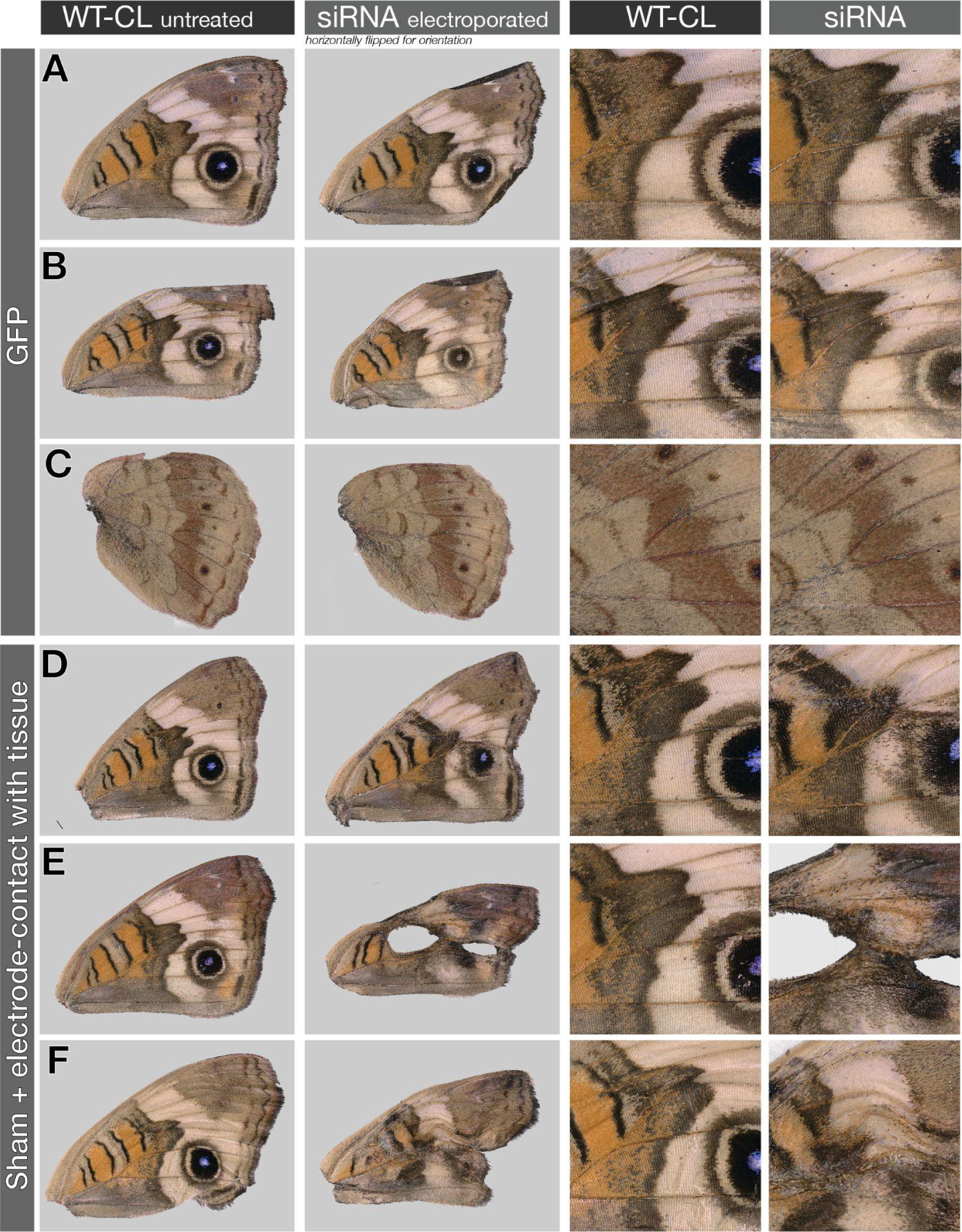
Pupal RNAi electroporation sham-negative controls and description of stress artefacts in *Junonia.* Examples of artefacts observed in sham electroporation procedures with GFP DsiRNA (A-B, ventral forewings; C, ventral hindwing), and following electroporation with the electrode directly in contact with the wing tissue (D-F). See Methods section for details.

**Figure S15.**
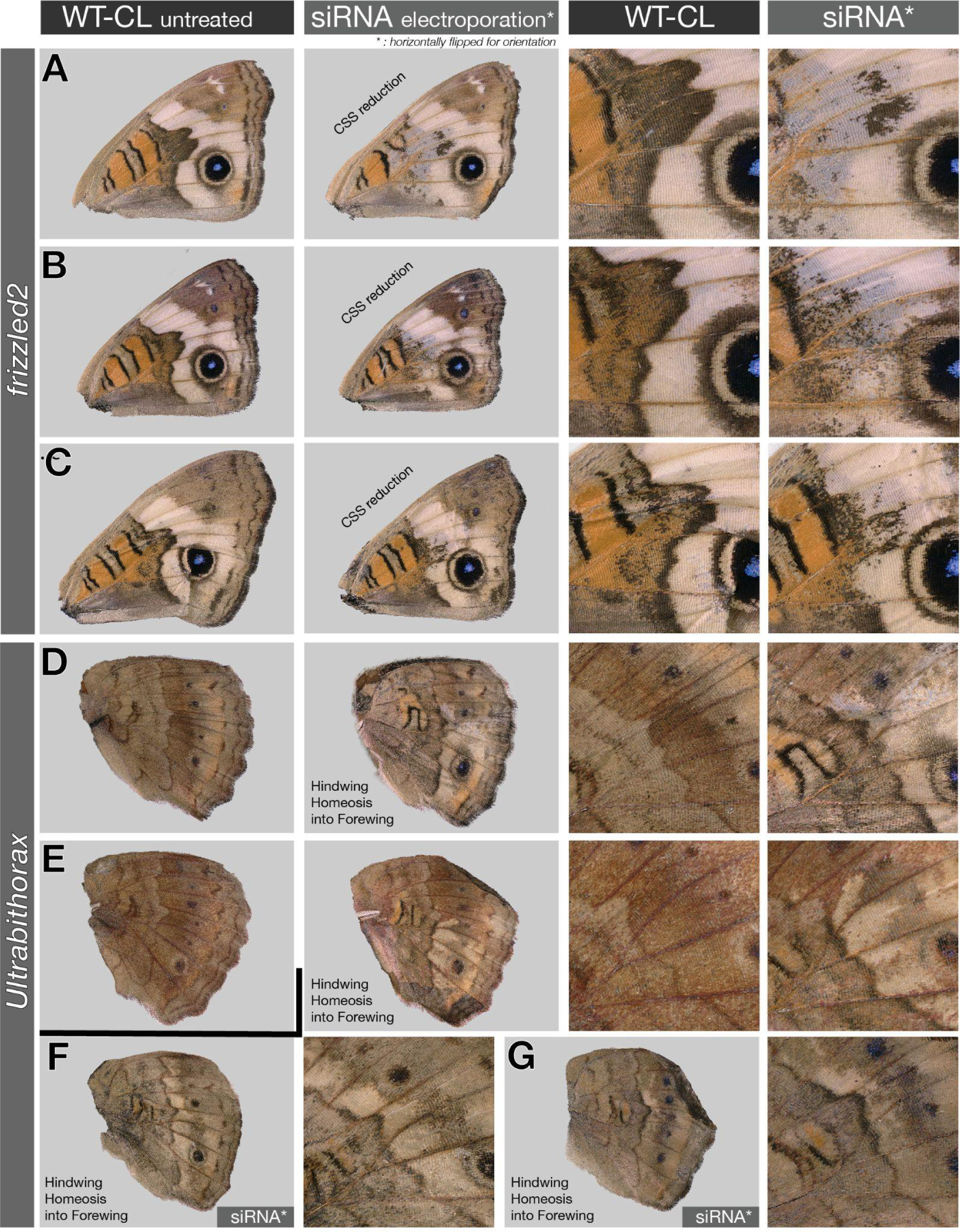
Pupal RNAi electroporation knockdowns of positive controls *fz2* and *Ubx* in *Junonia.* Representative examples of knockdown phenotypes obtained in forewings with *fz2* DsiRNA (A-C) and in hindwings with *Ubx* DsiRNA (D-G). Effects on CSS reduction (*fz2*) and partial homeoses of hindwings into forewings (*Ubx*) are consistent with expectations from CRISPR-induced KO phenotypes and validate our implementation of the RNAi electroporation method.

**Figure S16.**
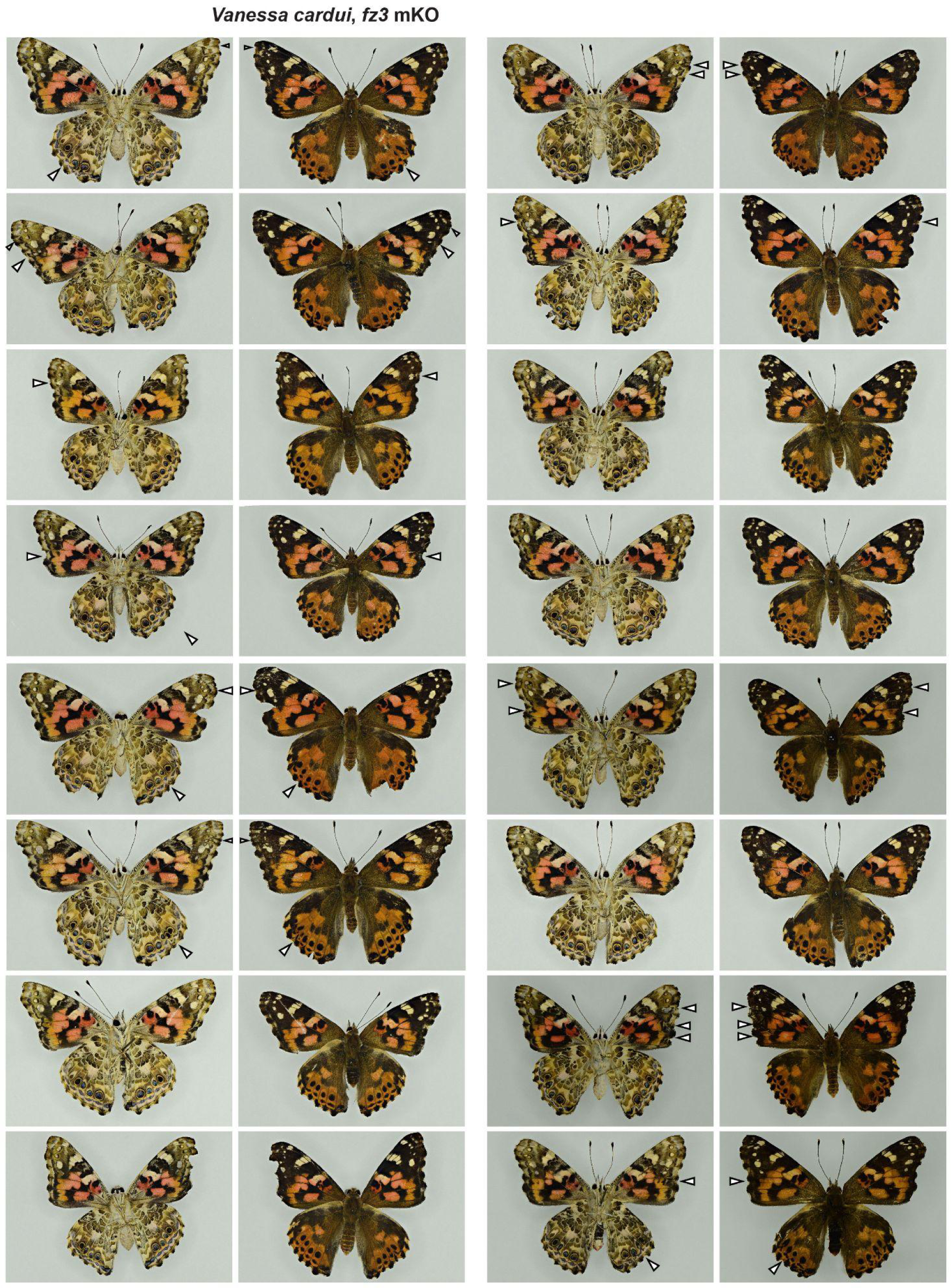
Representative *V. cardui fz3* crispant phenotypes. Ventral (left image) juxtaposed to dorsal (right image) sides for each individual. Arrowheads indicate ectopic veins.

**Figure S17.**
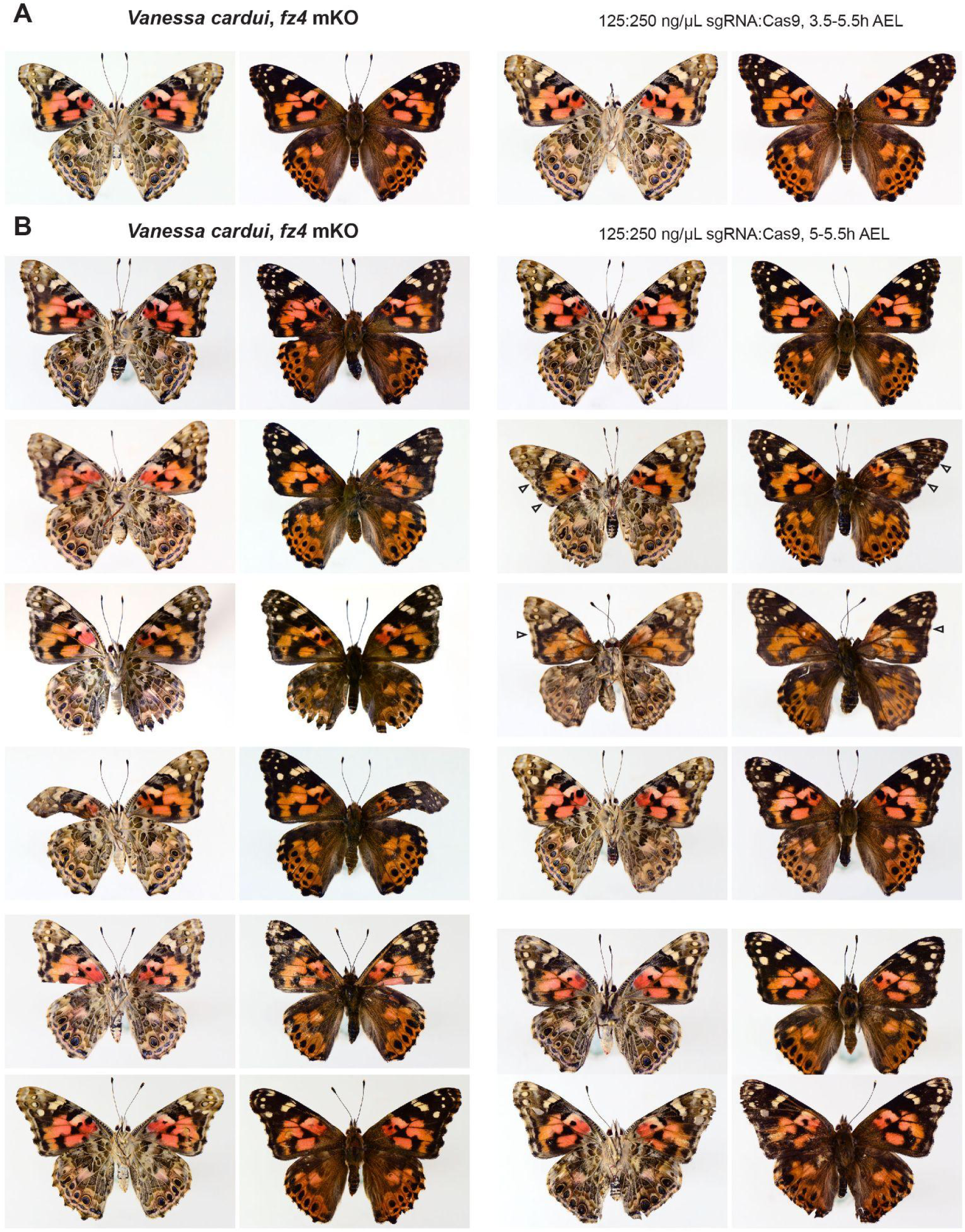
Representative *V. cardui fz4* crispant phenotypes. Ventral (left image) juxtaposed to dorsal (right image) sides for each individual.

**Figure S18.**
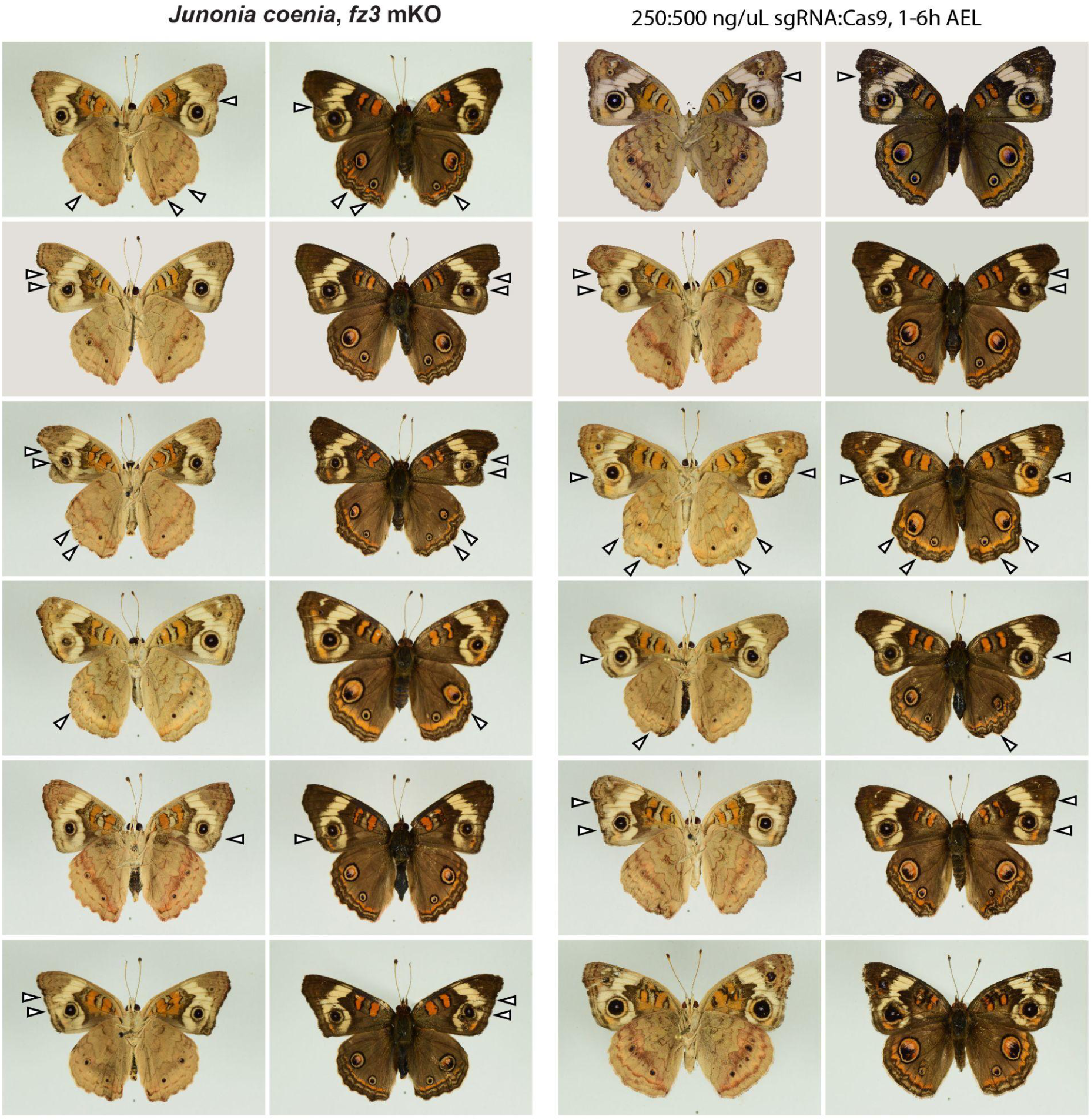
Representative *J. coenia fz3* crispant phenotypes. Ventral (left image) juxtaposed to dorsal (right image) sides for each individual. Arrowheads indicate ectopic veins.

**Figure S19.**
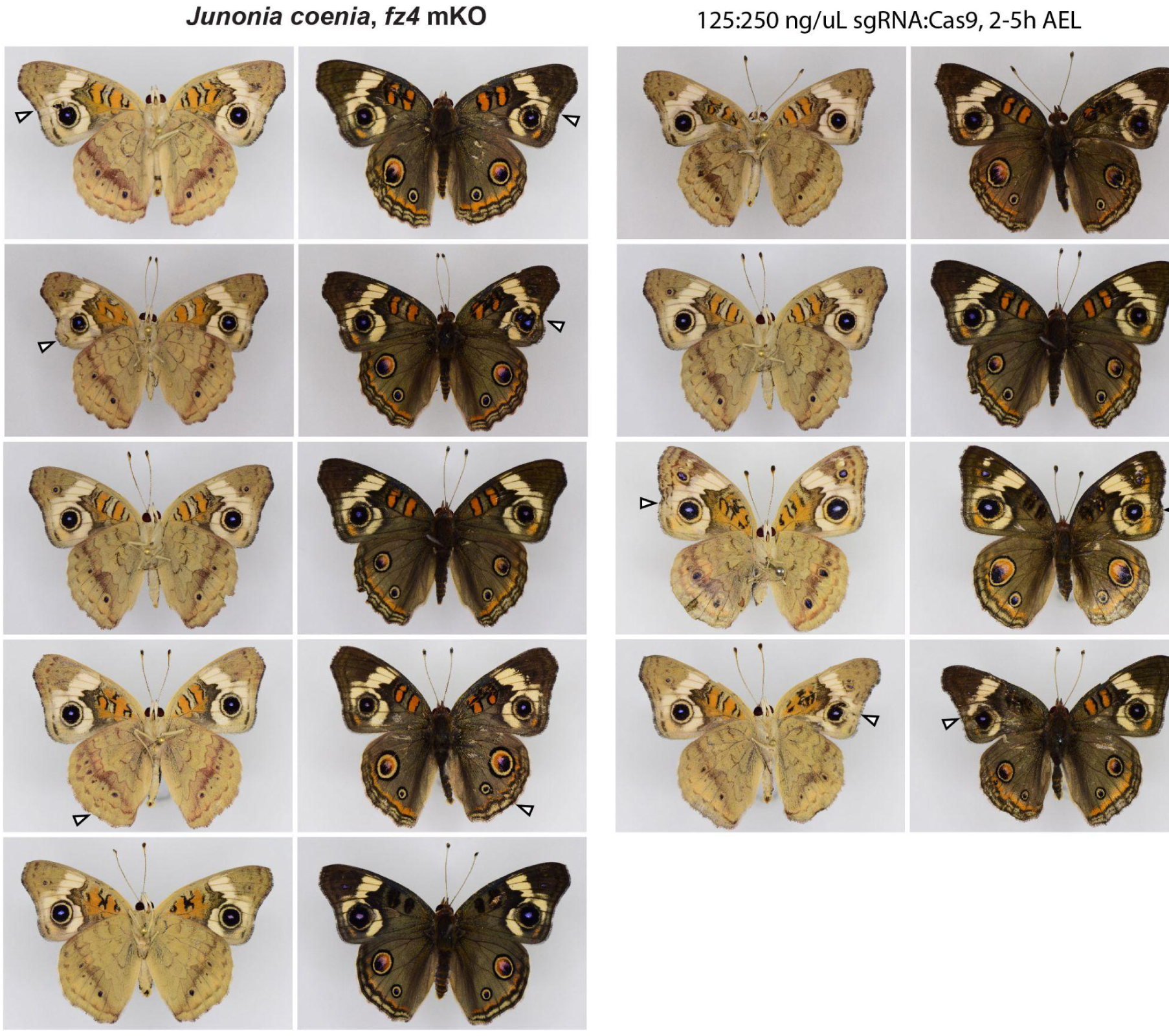
Representative *J. coenia fz4* crispant phenotypes. Ventral (left image) juxtaposed to dorsal (right image) sides for each individual.

**Figure S20.**
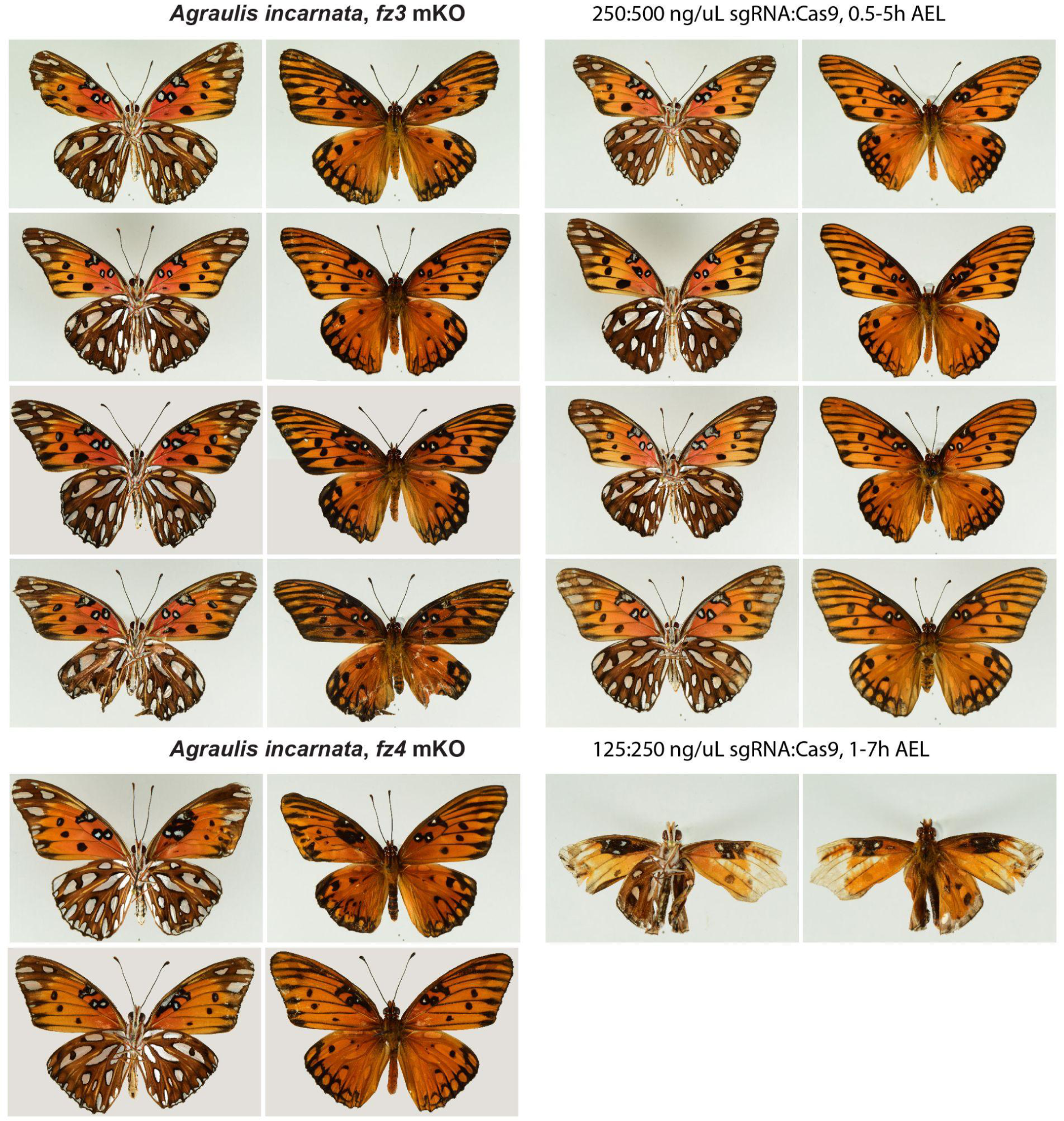
Representative *A. incarnata fz3* and *fz4* crispant phenotypes. Ventral (left image) juxtaposed to dorsal (right image) sides for each individual.

**Fig. S21.**
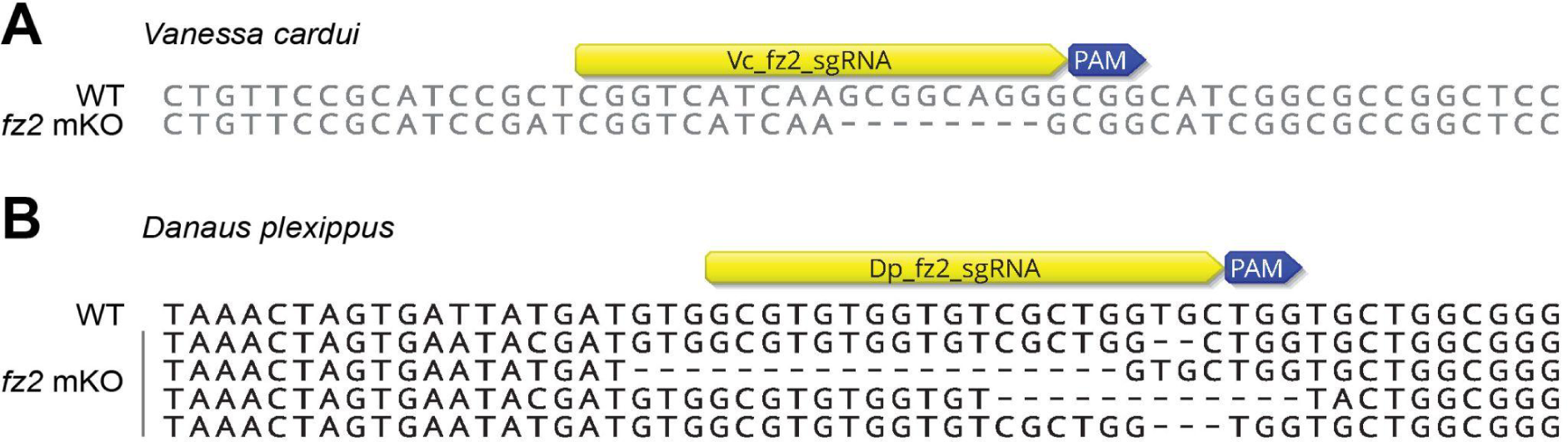
Genotyping of *fz2* G_0_ crispants. Thoracic muscle tissues were used for amplification of PCR products using *fz2*-specific primers (Table S3), and directly Sanger sequenced or isolated in single bacterial colonies by TA-cloning before sequencing. **(A)** Detection of a single frameshift mutation clone in a *V. cardui* crispant by direct Sanger sequencing. No wildtype sequence was recovered in this individual, implying the existence of a second mutant allele that was not amplified by PCR. DNA was directly amplified with the Phire Tissue Direct PCR Master Mix. **(B)** Detection of multiple frameshift mutations in two *D. plexippus* crispants

**Table S1.**
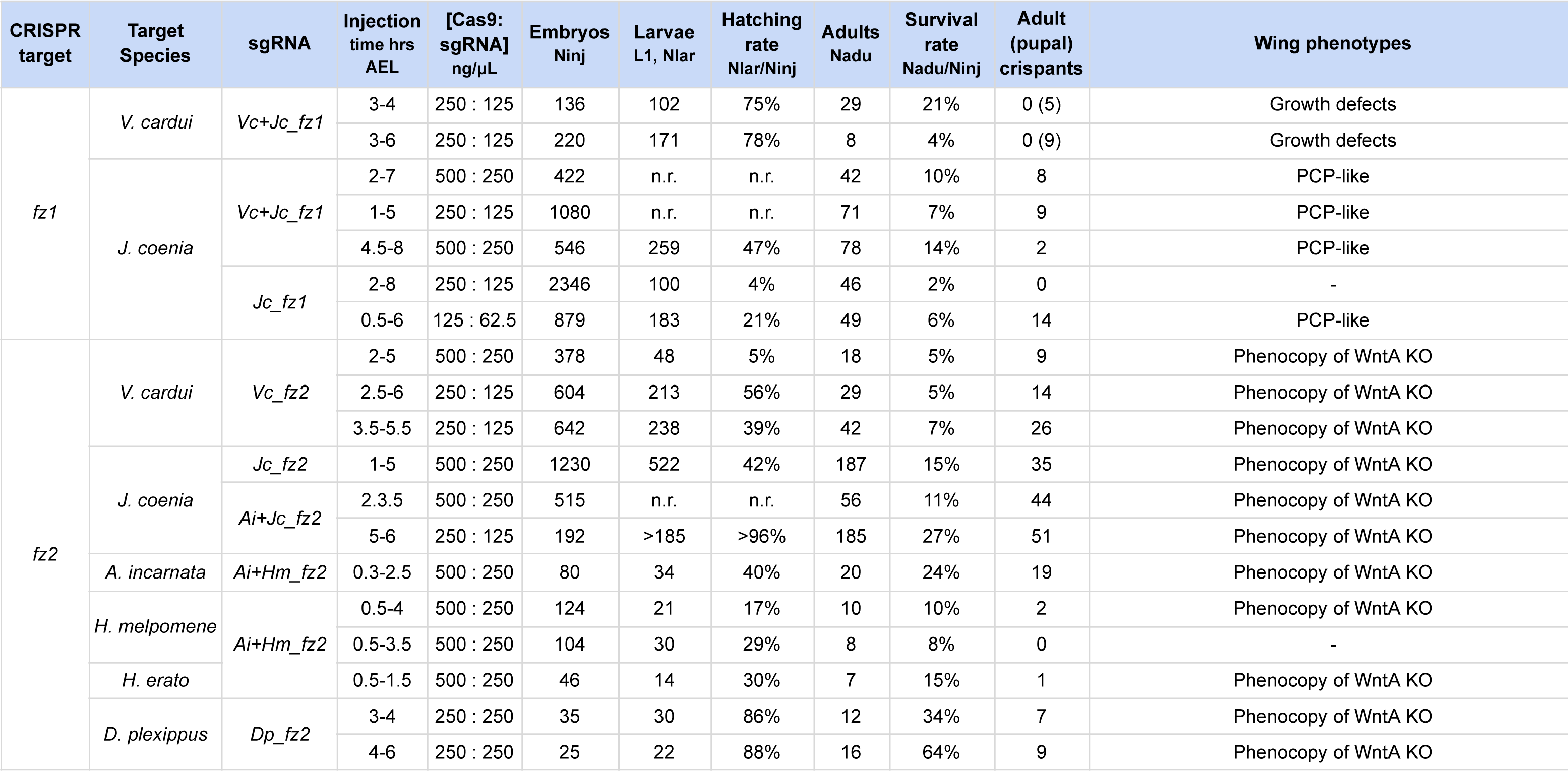
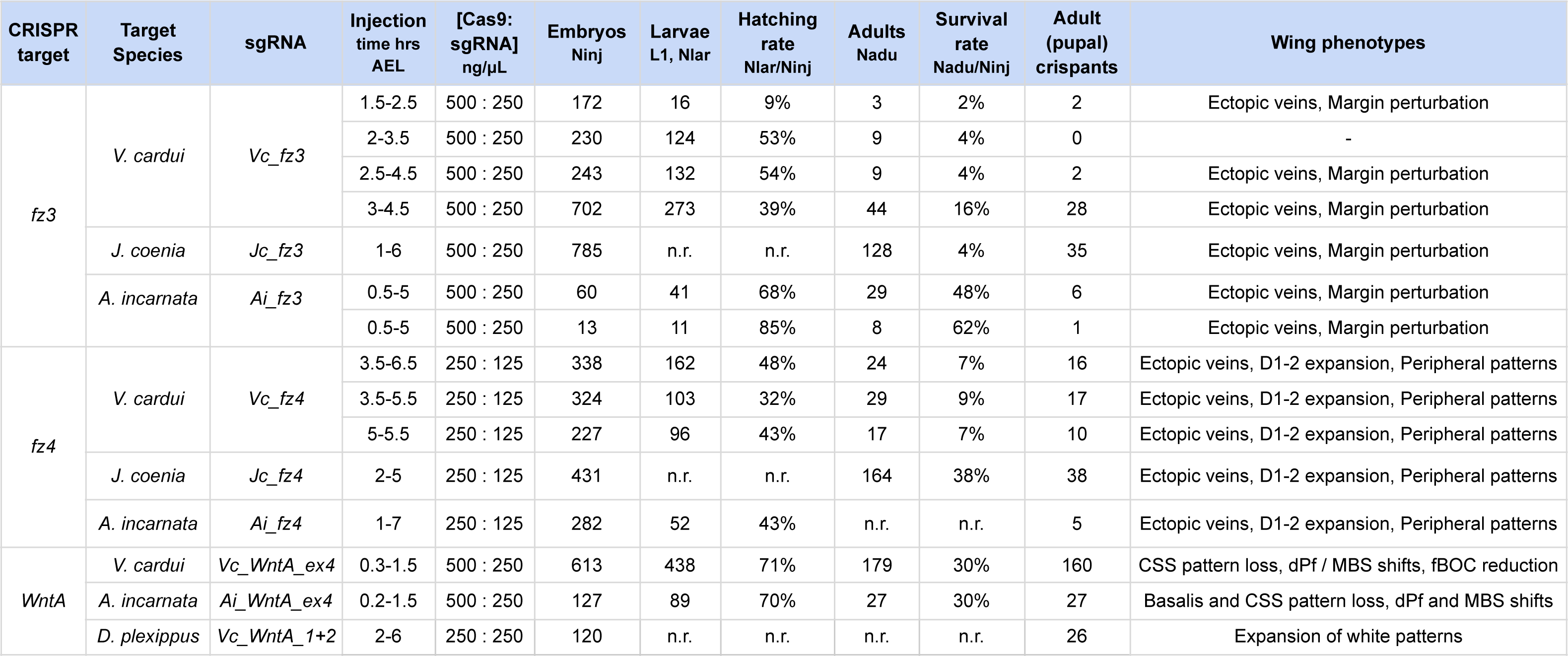
Summary of CRISPR-target mutagenesis experiments. n.r.: not recorded

**Table S2.**
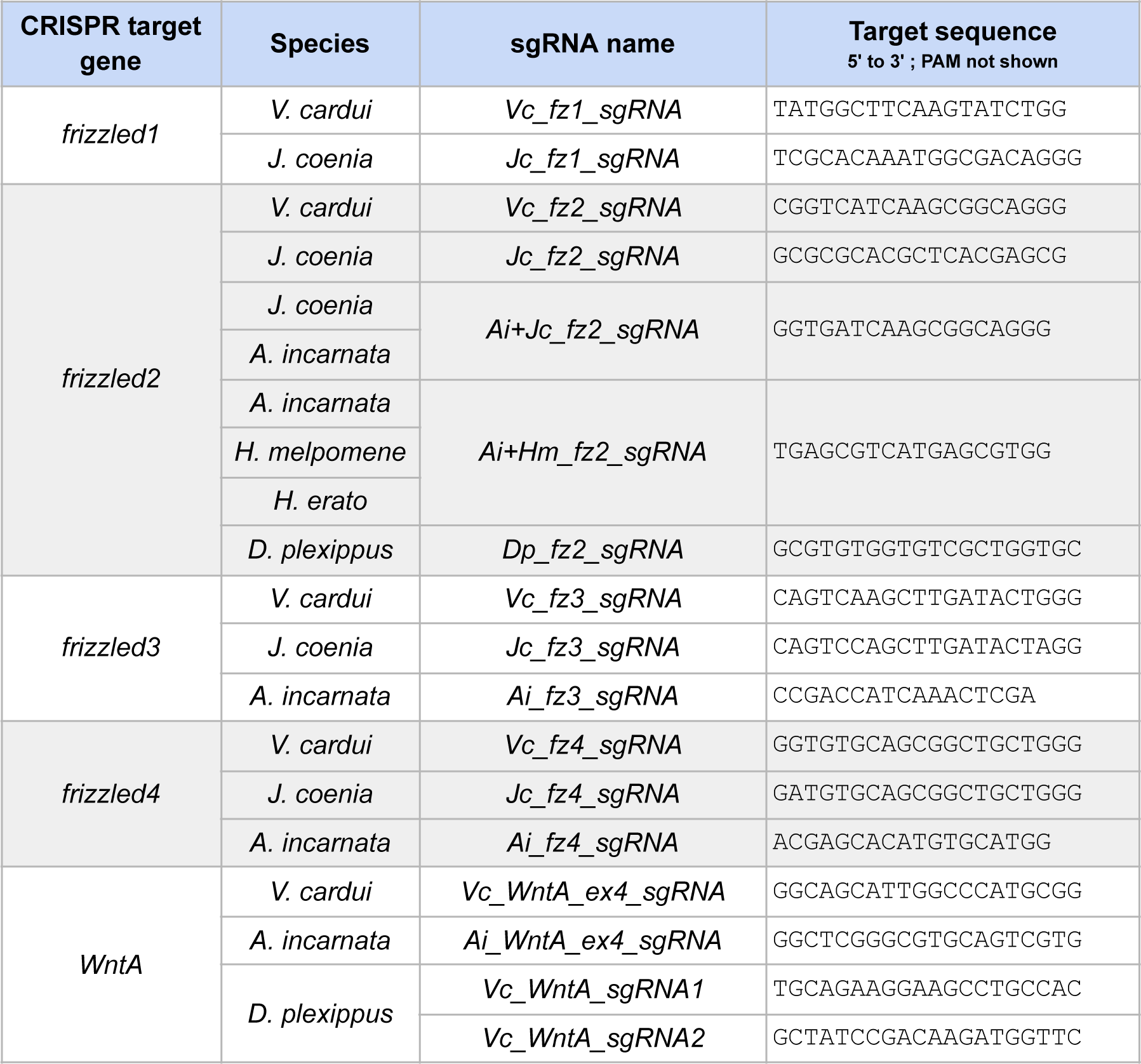
sgRNA sequences for CRISPR mosaic knock-outs.

**Table S3.**
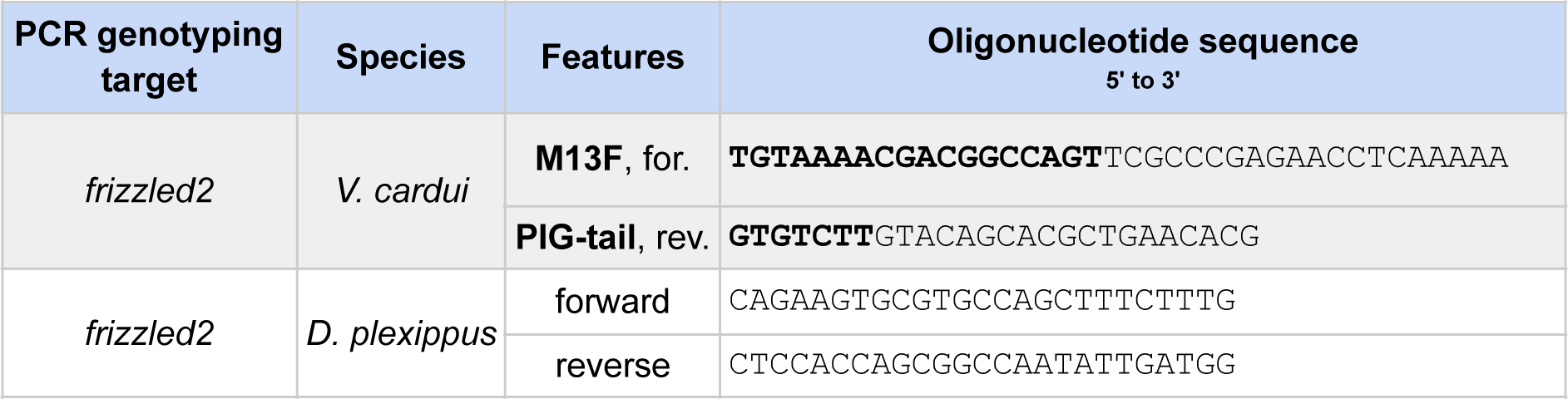
PCR primer sequences for the genotype confirmation of fz2 crispants.

**Table S4.**
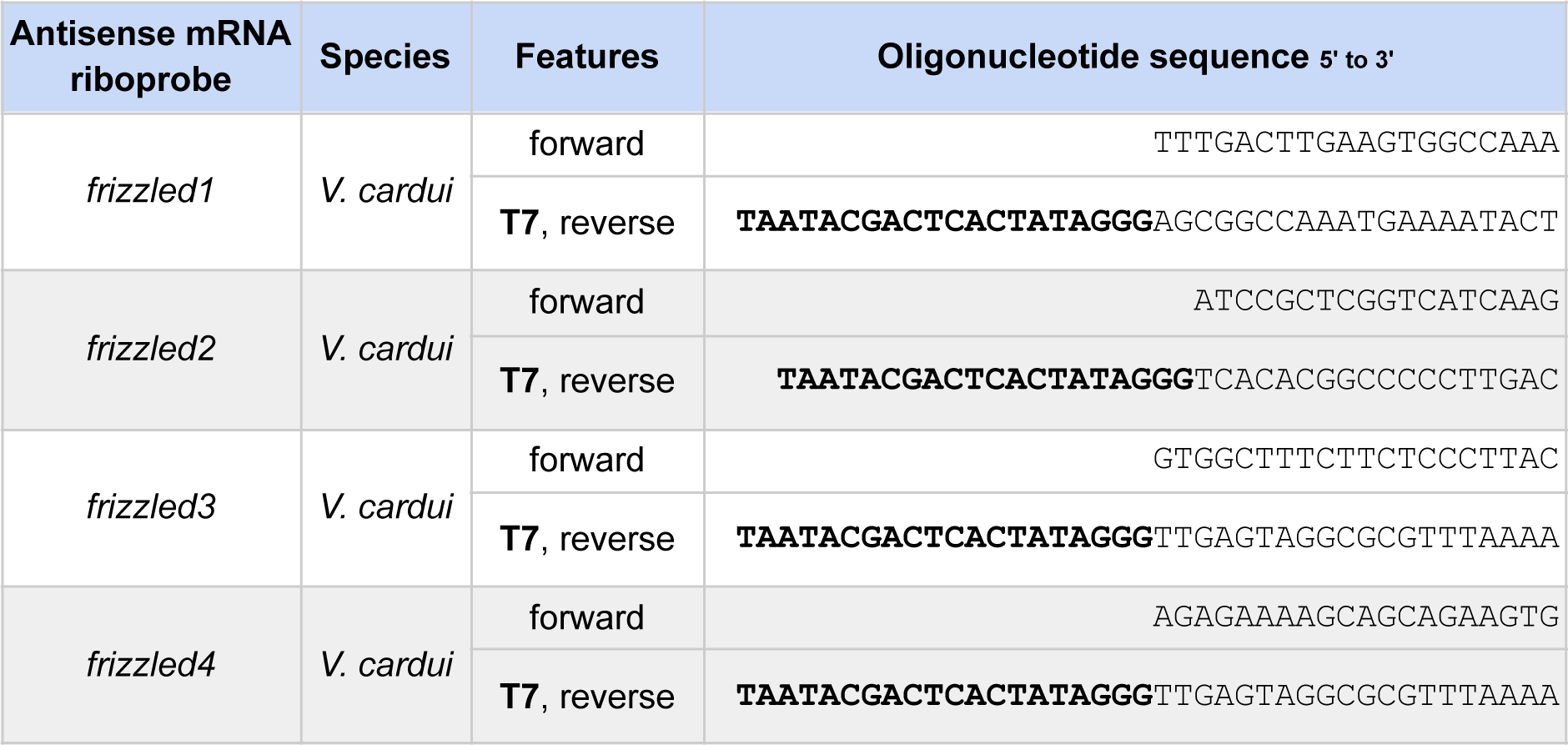
PCR primer sequences for the generation of riboprobe PCR transcription templates.

**Table S5.**
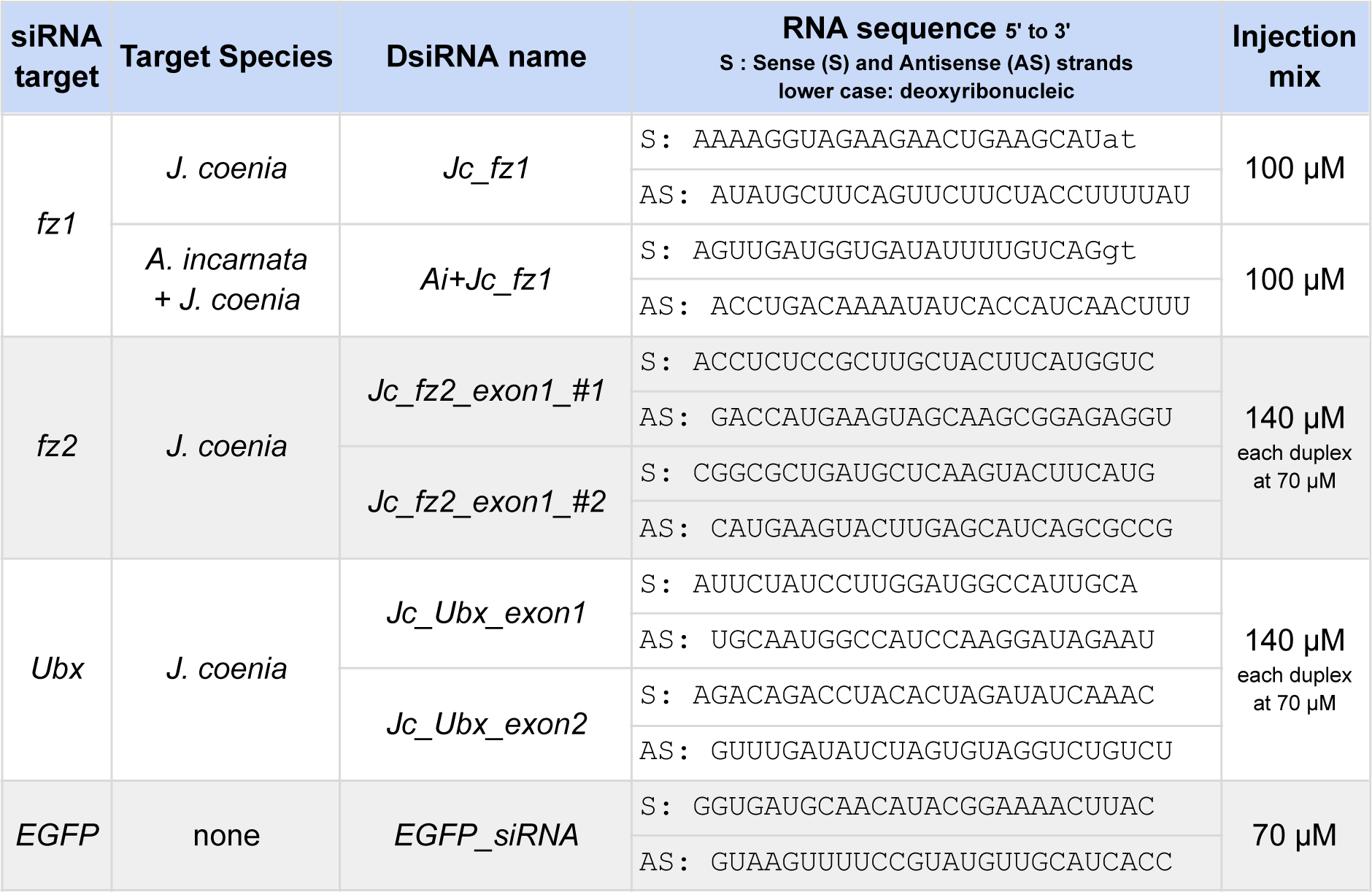
Dicer-substrate siRNA (DsiRNA) reagents used in pupal wing electroporations for gene expression knockdowns.

